# Zinc Finger Repressors mediate widespread *PRNP* lowering in the nonhuman primate brain and profoundly extend survival in prion disease mice

**DOI:** 10.1101/2025.03.05.636713

**Authors:** Shih-Wei Chou, Meredith A Mortberg, Kimberly Marlen, David S Ojala, Toufan Parman, Michael Howard, Kendrick DeSouza-Lenz, Yuan Lian, Mohadeseh Mehrabian, Matthew Tiffany, Annemarie Ledeboer, Patrick Dunn, Jing Hu, Kenia Guzman, Nikita Kamath, Morgan Cass, Samantha Graffam, Leo M H Xu, Vanessa Laversenne, Taylor L Corridon, Alaric Falcon, Garrett Lew, Satria Sajuthi, Lei Zhang, Giulia Cisbani, Finn Peters, Tim Fieblinger, Chiara Melis, Janica Barenberg, Marharyta Hnatiuk, Qi Yu, Daniel Chung, Mihika Jalan, Sarah Hinkley, Yanmei Lu, Kathleen Meyer, Alissa A Coffey, Amy M Pooler, Sonia M Vallabh, Eric Vallabh Minikel, Bryan Zeitler

## Abstract

Prion disease is a rapidly progressing and invariably fatal neurodegenerative disorder with no approved treatment. The disease is caused by the self-templated misfolding of the prion protein (PrP) into toxic species, ultimately leading to neurodegeneration and death. We evaluated a novel epigenetic regulation approach using Zinc Finger Repressors (ZFRs) to ablate PrP expression at the transcriptional level. When delivered using adeno-associated virus (AAV), ZFRs potently and specifically reduced prion mRNA expression by >95% *in vitro* and to near undetectable levels within single neurons *in vivo*. In wildtype mice, ZFRs stably lowered neuronal PrP expression throughout the central nervous system for at least 17 months. In mice inoculated with misfolded PrP, AAV-ZFRs given at either early or late disease stages profoundly extended lifespan, significantly reduced PrP in the brain, and improved an array of molecular, histological, biomarker, and behavioral readouts. Finally, we delivered a ZFR targeting the human prion gene (*PRNP)* to cynomolgus monkeys using a novel blood-brain-barrier penetrant AAV capsid. Extensive bulk and single-cell assessments revealed widespread ZFR expression and *PRNP* repression in all 35 brain regions assessed, providing the first demonstration of epigenetic regulation across the nonhuman primate neuraxis following a single intravenous (IV) dose. These results highlight the potential of a one-time IV administered ZFR treatment for prion disease and other neurological disorders.

## Introduction

Prion disease is a progressive, fatal neurodegenerative disorder caused by the misfolding of the normal cellular prion protein (PrP^C^) into a pathogenic “Scrapie” form (PrP^Sc^) (Prusiner 1982; Prusiner et al. 1998). Human prion disease can be categorized as sporadic (∼85%), genetic (∼15%), or acquired (<1%), with most cases being sporadic Creutzfeldt-Jakob disease (sCJD). Typically, sCJD patients have a median disease duration of around 5 months (Pocchiari et al. 2004). Symptoms may first present in cognitive, motor, or autonomic domains, but converge on profound dementia and akinetic mutism. Current diagnostic criteria include findings in brain magnetic resonance imaging (MRI) scans, Real-Time Quaking-Induced Conversion (RT-QuIC) for cerebrospinal fluid (CSF) PrP^Sc^, and electroencephalography (EEG) (Hermann et al. 2021). Common neuropathological hallmarks are neuronal loss, gliosis, and vacuolation in various brain regions (Takada and Geschwind 2013). Genetic prion disease, including familial CJD (gCJD), fatal familial insomnia (FFI), and Gerstmann–Straussler–Scheinker disease (GSS) are classified by genotypes, clinical symptoms, and neuropathological features. Although mutations in the human prion gene (*PRNP*) are dominant, the age of onset can be highly variable (Minikel et al. 2019), making it difficult to predict when pathogenic conversion will take place in the central nervous system (CNS). Worldwide, there are more than 5,000 new sCJD cases annually, with 1 in 6,000 deaths attributed to prion disease (Zerr et al. 2024; Maddox et al. 2020; Gao et al. 2024). Hence there is an urgent need to develop effective therapeutics.

Prion disease is well-modeled in animals, as many mammals are natural hosts of PrP^Sc^. In the laboratory, PrP^Sc^ injected into the rodent brain induces a robust, aggressive disease course that mimics many hallmarks of human prion disease, including biomarker, histopathological, and behavioral changes. In mice, the most commonly studied laboratory prion strain, Rocky Mountain Laboratory (RML), induces a rise in plasma neurofilament light chain (NfL) levels at ∼60 days post-inoculation (dpi), onset of clinical symptoms at ∼120 dpi, and mortality at ∼170 dpi (Minikel et al. 2020).

While there is currently no approved treatment for prion disease, the histopathological, biochemical, and genetic evidence all converge on PrP^C^ as the primary drug target for treating prion disease (Prusiner 1982; Vallabh et al. 2020). Prion gene knockout (KO) animals are completely resistant to prion infection (Bueler et al. 1993; Sailer et al. 1994; Salvesen et al. 2020). Reducing PrP^C^ expression to 50% in heterozygous genetic KO mice or by chronic treatment with PrP-lowering antisense oligonucleotides (ASOs) can extend the survival of PrP^Sc^-infected mice (Bueler et al. 1994; Raymond et al. 2019). Nonetheless, all of these PrP^Sc^-infected mice ultimately succumb to terminal disease, indicating that halting disease progression will require more than 50% knockdown. Interestingly, prion neurotoxicity is cell autonomous, meaning that any neuron not expressing PrP^C^ is protected, even in the presence of misfolded PrP^Sc^ from neighboring cells (Brandner, Isenmann, et al. 1996; Lakkaraju et al. 2022). Due to a lack of tools capable of achieving complete PrP knockdown within individual neurons *in vivo*, it is currently unknown whether 100% pharmacologic lowering of PrP^C^ in neurons could halt or prevent disease progression.

Although our understanding of normal PrP^C^ function is incomplete, evidence from KO mice, goats, sheep, and cattle suggest it is not essential for normal development, function, or survival (Bueler et al. 1992; Yu et al. 2009; Richt et al. 2007; Benestad et al. 2012). No significant CNS deficits have been observed in adult animals with post-developmental genetic knockout of PrP^C^, suggesting that PrP^C^ lowering in the brain is a viable therapeutic approach for prion disease (Mallucci et al. 2003). Most reported phenotypes in KO mice have not been replicated, and many have been refuted or assigned to genetic artifacts (Wulf, Senatore, and Aguzzi 2017). A demyelinating neuropathy has been associated with PrP KO in mice (Bremer et al. 2010) but is phenotypically mild and observed only in the peripheral nervous system of aged homozygous knockouts. Repeated intracerebroventricular (ICV) administration of an ASO targeting mouse *Prnp* can extend survival of RML-inoculated mice (Raymond et al. 2019; Minikel et al. 2020), and an intrathecally administered ASO is currently in a Phase I clinical trial for prion disease (NCT06153966). Nevertheless, repetitive ASO dosing maximally achieves ∼50% knockdown of *Prnp* mRNA expression, without complete knockdown in any individual cell, and with only limited coverage of deeper brain regions (Jafar-Nejad et al. 2021; Mortberg et al. 2023). Developing additional therapeutic modalities that can achieve greater bulk and single-cell knockdown, more uniform CNS distribution, and a less burdensome administration regimen remains an important goal.

One potential innovative treatment of prion disease is to use AAV delivery of an epigenetic regulator to achieve sustained *PRNP* transcriptional repression in the brain. Epigenetic regulators have been used to suppress the expression of various genes linked to neurological disorders in preclinical models for conditions such as chronic pain, Huntington’s disease, and Alzheimer’s disease (Samie et al. 2024; Zeitler et al. 2019; Wegmann et al. 2021; Moreno et al. 2021; Tanenhaus et al. 2022). Zinc Finger Proteins (ZFPs) are particularly well-suited for this purpose, as they are naturally occurring transcription factors with high DNA-binding specificity that have evolved to potently regulate eukaryotic gene expression. Zinc Finger Repressors (ZFRs), which consist of an engineered Zinc Finger DNA-binding array and a human Krüppel-associated box (KRAB) transcriptional repression domain, offer several potential advantages over other genomic medicine approaches for modulating gene expression. ZFRs are compact, enabling unrestricted packaging into AAV vectors, and present a potentially lower risk of immunogenicity compared to bacterially derived CRISPR-based tools. Because ZFRs reversibly recruit endogenous epigenetic silencing complexes, they do not permanently alter or damage the genome or epigenome. As neurons and glial cells are post-mitotic, cells transduced with prion gene-targeted ZFRs are expected to have lifelong ZFR expression and concomitant *PRNP* knockdown, avoiding peak-trough differences in drug levels and the need for repeat dosing.

A major challenge in developing ZFR-based therapies for human CNS diseases has been the lack of AAV vectors capable of achieving widespread brain distribution in primates. Naturally occurring serotypes such as AAV9 exhibit limited, heterogeneous, or focal biodistribution when administered via CSF or intraparenchymal routes (Hudry and Vandenberghe 2019; Meseck et al. 2022). For prion disease, which affects the entire brain, widespread neuraxis transduction by a blood-brain-barrier (BBB) penetrant AAV capsid may provide the greatest potential for clinical benefit. Recent progress in capsid engineering has led to the discovery of novel BBB-penetrant capsids that mediate enhanced CNS delivery in mice, humanized mouse models, and nonhuman primates (NHPs), each with translational potential for delivery in humans (Shay et al. 2023; Huang et al. 2024; Moyer et al. 2024; Tiffany et al. 2024). Such efforts have yielded potential delivery solutions for a ZFR-based epigenetic regulator via a minimally invasive, one-time IV administration. However, despite this important progress, it is unknown whether these novel capsids can achieve sufficient delivery of an epigenetic regulator in the NHP brain for any CNS disease target.

Here, we engineered ZFRs to specifically repress PrP^C^ expression in the brain and evaluated their performance in mice and NHPs across a range of molecular, cellular, and functional endpoints. We tested different promoters for cell-type specific or ubiquitous expression using bulk tissue and single-cell measurements. We demonstrate that a one-time IV administration of AAV-delivered prion-targeted ZFRs to mice with prion disease specifically lowered brain *Prnp* mRNA and PrP protein, increased median survival time, delayed plasma NfL increases, and arrested the neuropathological changes associated with prion disease. Finally, we demonstrate for the first time that delivery of a ZFR to NHPs using an engineered BBB-penetrant AAV capsid can achieve widespread transduction and *PRNP* repression at levels associated with profound survival benefits in prion-infected mice.

## Results

### ZFRs potently repress mouse *Prnp* with high specificity *in vitro*

We designed mZFRs to specifically target the transcriptional regulatory region of the mouse *Prnp* gene and screened these for repression of *Prnp* mRNA in mouse Neuro2A (N2A) cells. Reverse transcription-quantitative polymerase chain reaction (RT-qPCR) analysis of N2A cells after mZFR nucleofection identified dozens of highly potent ZFRs that reduced *Prnp* mRNA by 50-99% (Figure 1A). When packaged into AAV6 for optimal transduction in vitro (Ellis et al. 2013), two lead mZFRs (mZFR1 and mZFR2) expressed using a human synapsin 1 (hSYN1) promoter potently reduced *Prnp* mRNA levels in primary mouse cortical neuron cultures in a dose-dependent fashion (Figure 1B). Whole-transcriptome profiling of >20,000 transcripts revealed no off-target genes were significantly downregulated for the selected mZFRs (Figure 1C). In contrast, a less-specific ZFR (mZFR3) associated with up- and downregulation of several off-target genes was not advanced for further evaluation (Supplementary Figure 1).

**Figure 1.**
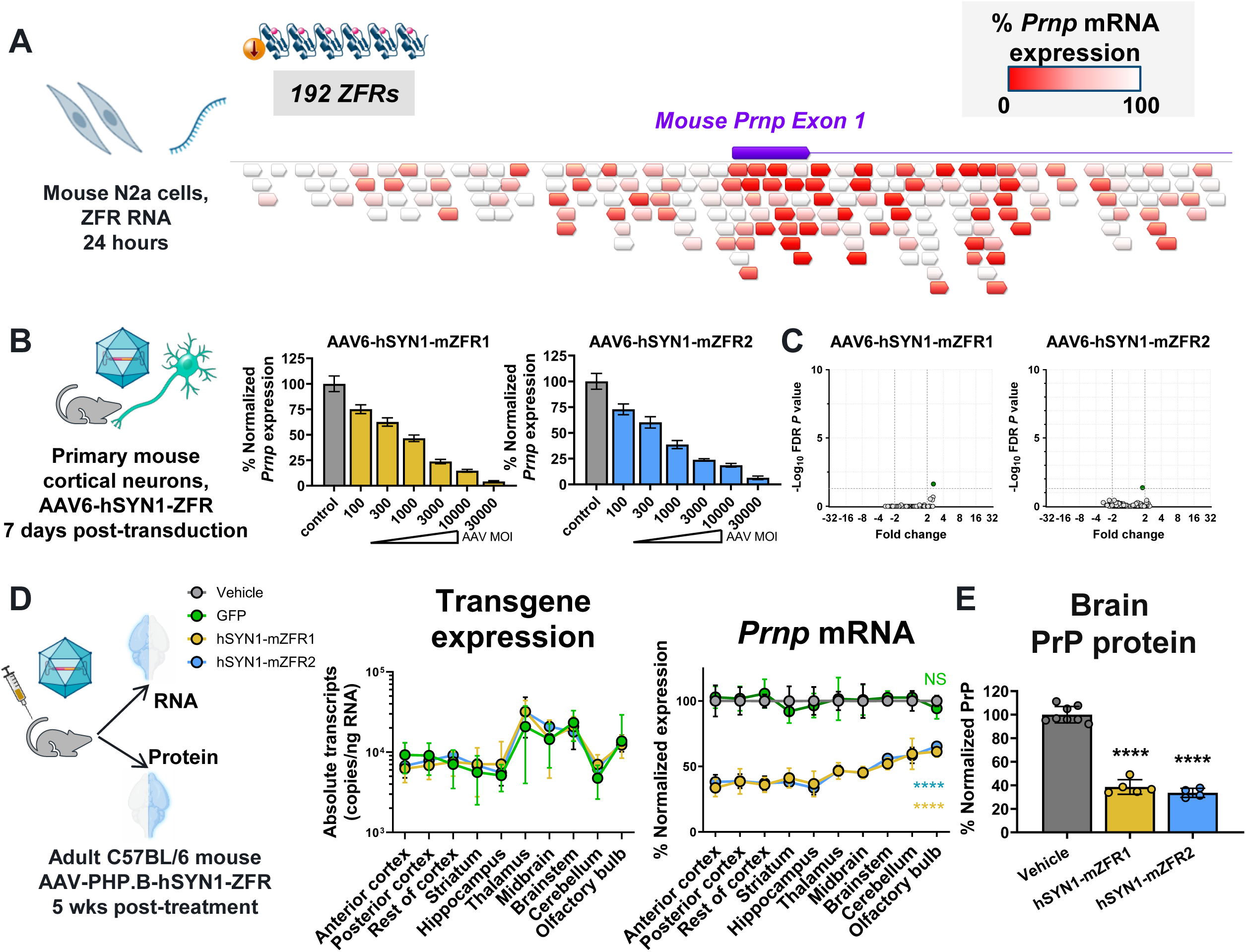
Characterization of potent and specific mZFRs targeting mouse *Prnp in vitro* and in vivo. **(A)** 192 mZFRs designed to target various locations in the mouse *Prnp* gene were tested in Neuro-2A cells with ∼30% of mZFRs achieving at least 50% repression of the *Prnp* transcript at 1000 ng dose. **(B)** On-target analysis in primary mouse cortical neuron (MCN) culture. Percent normalized *Prnp* expression was measured 7 days after mZFR transduction and scaled to control levels (mean ± standard deviation (SD)). **(C)** Off-target analysis of AAV6-hSYN1-mZFRs in MCNs dosed at 3E+3 Multiplicity Of Infection (MOI) were evaluated. *P*-values were calculated based on False Discovery Rate (FDR) value < 0.05. **(D)** Brain hemispheres collected from mice 5 weeks after dosing with vehicle (n=8 animals, grey), GFP (n=6, green), hSYN1-mZFR1 (n=5, yellow), and hSYN1-mZFR2 (n=4, blue) were micro-dissected into ten regions. RNA extracted from each region was analyzed by RT-qPCR. Transgene (mean ± SD, absolute copies/ng RNA input) and normalized *Prnp* expression (mean ± SD, scaled to the mean of vehicle group). Two-way ANOVA with Dunnett’s multiple comparisons test, **** P < 0.0001. **(E)** Normalized PrP protein expression level (mean ± SD, scaled to the mean of vehicle group) measured by ELISA. One-way ANOVA with Dunnett’s multiple comparisons test, **** P < 0.0001.

### ZFR-mediated reduction of *Prnp* mRNA and PrP protein in the mouse brain

We first evaluated the performance of mZFR1 and mZFR2 in wildtype C57BL/6 mice. We selected the PHP.B capsid for these studies, which efficiently crosses the BBB in this mouse strain following IV administration (Deverman et al. 2016). The AAV constructs included a hSYN1 promoter to limit CNS expression to neurons (Wegmann et al. 2021). Adult mice received either AAV-PHP.B-hSYN1-mZFR1 (hSYN1-mZFR1) or AAV-PHP.B-hSYN1-mZFR2 (hSYN1-mZFR2) at 1E+14 vg/kg. Brain tissues were collected for RT-qPCR and enzyme-linked immunosorbent assay (ELISA) analysis 5 weeks after dosing. Treatment with hSYN1-mZFRs led to brain-wide *Prnp* mRNA reduction of 34∼66% when compared with the vehicle control group (Figure 1D). Moreover, animals that received a negative control vector, AAV-PHP.B encoding a green fluorescent protein (GFP) transgene, did not show any *Prnp* repression in the brain, indicating the target specificity of hSYN1-mZFRs in vivo (Figure 1D). When compared to the vehicle-treated group, the average PrP protein levels in the full brain hemisphere were also significantly reduced by 61 and 66% for hSYN-mZFR1 and hSYN1-mZFR2, respectively (Figure 1E).

### Cell Types Involved in Brain *Prnp* Expression

Next, the contribution of different cell types to brain *Prnp* expression was assessed in a promoter comparison study using mZFR1 under the control of one of three promoters with distinct expression patterns: hSYN1 (pan-neuronal), GfaABC1D (astrocytic), or CMV (ubiquitous) (Thiel, Greengard, and Sudhof 1991; Lee et al. 2008). To facilitate detection of transduced cells, each test article was constructed as AAV-PHP.B-Promoter-mZFR1-T2A-GFP-H2B, where T2A is a ribosomal skipping sequence separating mZFR1 and GFP, and a histone 2B (H2B) sequence was fused to GFP to confine its distribution to the nucleus. The construct and study design are shown in Figure 2A. Treatment with these constructs led to significant repression of *Prnp* mRNA across all brain regions and spinal cord for all promoter types when compared to the control group (Figure 2B, left panel). The hSYN1 construct was highly selective for the brain, with no statistically significant *Prnp* repression detected in peripheral tissues compared to the vehicle group. The GfaABC1D promoter was similarly inactive in most peripheral tissues; however, it also resulted in less *Prnp* reduction across all CNS regions compared to hSYN1. The CMV promoter resulted in potent *Prnp* repression in several CNS regions, as well as in liver, heart, kidney, and spleen, indicating functional ZFR expression in non-neuronal cells (Figure 2B, right panel). PrP protein levels from brain hemispheres were analyzed by ELISA. While all three constructs displayed a statistically significant lower level of PrP than controls, animals treated with the hSYN1 construct showed the greatest reduction (∼56%) compared with animals treated with constructs containing GfaABC1D (∼43%) or CMV (∼43%) promoter (Figure 2C). Similarly, mean % reduction in PrP protein levels in CSF was 68%, 46%, and 74% for hSYN1-, GfaABC1D-, and CMV-containing constructs, respectively (Figure 2D).

**Figure 2.**
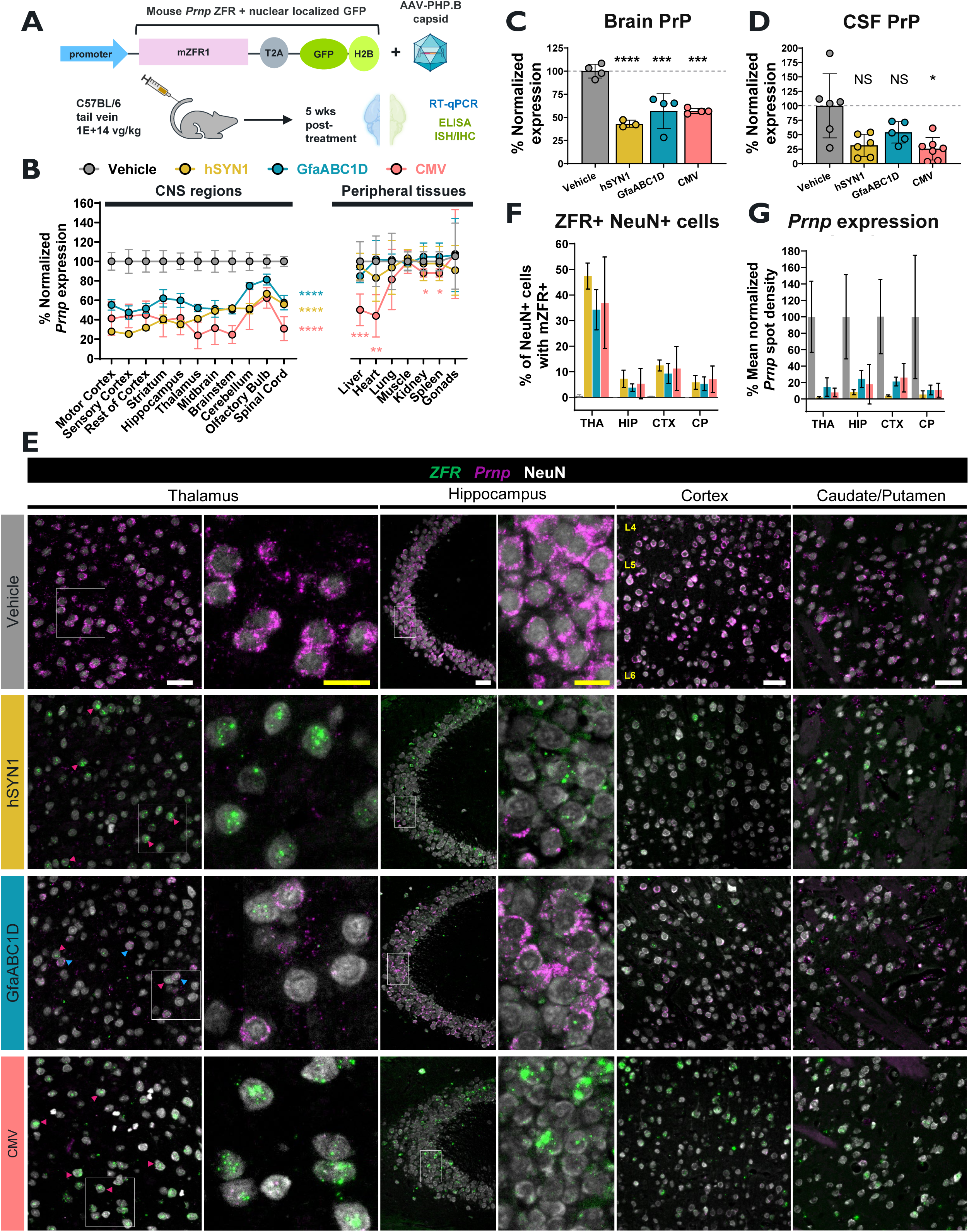
*Prnp* repression in mouse brain with mZFR1 driven by different promoters at the bulk and single-cell level. **(A)** Schematic diagram of the construct and experimental design. **(B)** RT-qPCR analysis of *Prnp* expression (% normalized values were scaled to the mean of the vehicle group, mean ± SD) in 11 CNS regions (brain and spinal cord) and 7 peripheral tissues. For the brain regions and spinal cord, all three promoter-mZFR1 groups showed statistically significant *Prnp* repression (****) compared to the vehicle control. For peripheral tissues, only the CMV-mZFR1-T2A-GFP-H2B (CMV) group resulted in statistically significant reduction of *Prnp* seen in liver (***), heart (**), kidney (*), and spleen (*) (two-way ANOVA test with Dunnett’s multiple comparisons test). **(C, D)** PrP protein reduction observed in brain and CSF collected from different groups of mZFR1-dosed mice. **(C)** Normalized PrP protein expression level analyzed by ELISA in brain hemispheres (mean ± SD). One-way ANOVA test with Dunnett’s comparison between vehicle and individual treated group. P-value: <0.0001**** for hSYN1, 0.0005*** for GfaABC1D and CMV. **(D)** Normalized PrP protein expression level analyzed by Meso Scale Discovery (MSD) in CSF (mean ± SD). Kruskal-Wallis test with Dunnett’s comparison between vehicle and individual treated group. P-value: 0.06 (hSYN1), >0.99 (GfaABC1D), 0.01* (CMV). **(E)** Representative microscopy images from coronal brain slices for thalamus (THA), hippocampus (HIP), cortex (CTX), and caudate/putamen (CP). White and yellow scale bars are 50 and 20 µm, respectively. Blue arrows, cells with detectable level of *Prnp* expression. Pink arrows, cells transduced with ZFR and showed low to no *Prnp* expression. L4, L5, and L6 indicate cortical layers 4, 5, and 6, respectively. **(F)** Percentage of ZFR-positive neurons in brain regions for vehicle and treatd groups, mean ± SD for n = 4 animals, except hSYN1 group (n = 3). No significant difference was observed between the three different promotors for a given brain region (unpaired t-test comparing two groups each from a given brain region). **(G)** *Prnp* mean normalized spot density in all NeuN+ (vehicle) or ZFR+ NeuN+ (treated groups) cells (mean ± SD). Two-way ANOVA using Tukey’s multiple comparison test demonstrated that *Prnp* reduction from all three promoter-mZFR1-treated groups were significant in four brain regions analyzed. There were no significant differences among treatment groups.

To assess promoter cell-type specificity and selectivity, coronal brain sections were stained using in situ hybridization (ISH) probes for mZFR1 and mouse *Prnp* combined with immunohistochemistry (IHC) using cell-type specific antibodies for NeuN (neuronal marker) and S100ß (astrocyte marker) (Figure 2E-G). Expression of mZFR1 was found throughout the brain in NeuN+ cells in the hSYN1 and CMV groups but was less evident for the GfaABC1D group (Figure 2E). Despite the differences observed in bulk RT-qPCR measurements, there were no significant differences in the percentage of mZFR+ NeuN+ cells among the three constructs. This may be due to the cross-reactivity of the mZFR1 ISH probe for both RNA and vg DNA, thereby also detecting cells that are transduced but not expressing mZFR1. Because of this – and the presence of neuronal progenitor cells in the analyzed brain regions that can be labeled by both NeuN and S100ß antibodies (Supplementary Figure 2.1) – it is unclear which constructs promote efficient transgene expression in astrocytes. Critically, *Prnp* staining demonstrated that neurons transduced with either promoter construct had significantly lower or no detectable *Prnp* mRNA compared to the vehicle control group (Figure 2E). Expression levels of *Prnp* mRNA transcripts in neurons were quantified as mean spot density from NeuN+/mZFR+ cells in the treated groups and normalized against the mean spot density from NeuN+ cells of the vehicle control group for each brain region and animal. Greater than 90% reduction of *Prnp* mRNA level was observed in neurons within the thalamus, hippocampus, cortex, and caudate/putamen of animals that received the hSYN1 promoter construct when compared to the vehicle control group (Figure 2G). From the same brain regions analyzed, neuronal *Prnp* mRNA reduction of 75-80% was observed for GfaABC1D and CMV constructs. The correlation between mZFR1 expression and decreased *Prnp* expression was nonlinear, i.e. cells with more mZFR1 transcripts did not necessarily lead to lower *Prnp* expression than those with fewer mZFR1 transcripts; the presence of even the lowest level of mZFR1 transcripts in a given cell appears sufficient to significantly reduce *Prnp* in that cell. Interestingly, we also observed *Prnp* repression in many cells without detectable mZFR1-signal, suggesting the threshold of mZFR1 expression required to silence *Prnp* is below the limit of detection of this assay. The number of NeuN+/mZFR1+ cells is therefore likely to underestimate the total number of transduced cells in the brain.

To further distinguish cell types expressing mZFR1 and *Prnp* we conducted single nucleus transcriptome sequencing of the visual cortex. Cell type clustering provided a comprehensive landscape of target gene repression in various cell types (Figure 3A-B). Despite AAV-PHP.B transducing a majority of neurons and a high percentage of astrocytes (Deverman et al. 2016), the transgene transcript was detected in a relatively small number of cells for all constructs (Figure 3C). This is expected because single nucleus sequencing is relatively sparse (i.e., not all expressed transcripts in all cells yield even one sequencing read), and biased towards transcripts that are significantly larger than mZFR1 (Chamberlin et al. 2024). One interpretation is that cells with a detected mZFR1 transgene sequencing read (ZFR+) were definitely transduced, while those without (ZFR-) include both transduced and untransduced cells.

**Figure 3.**
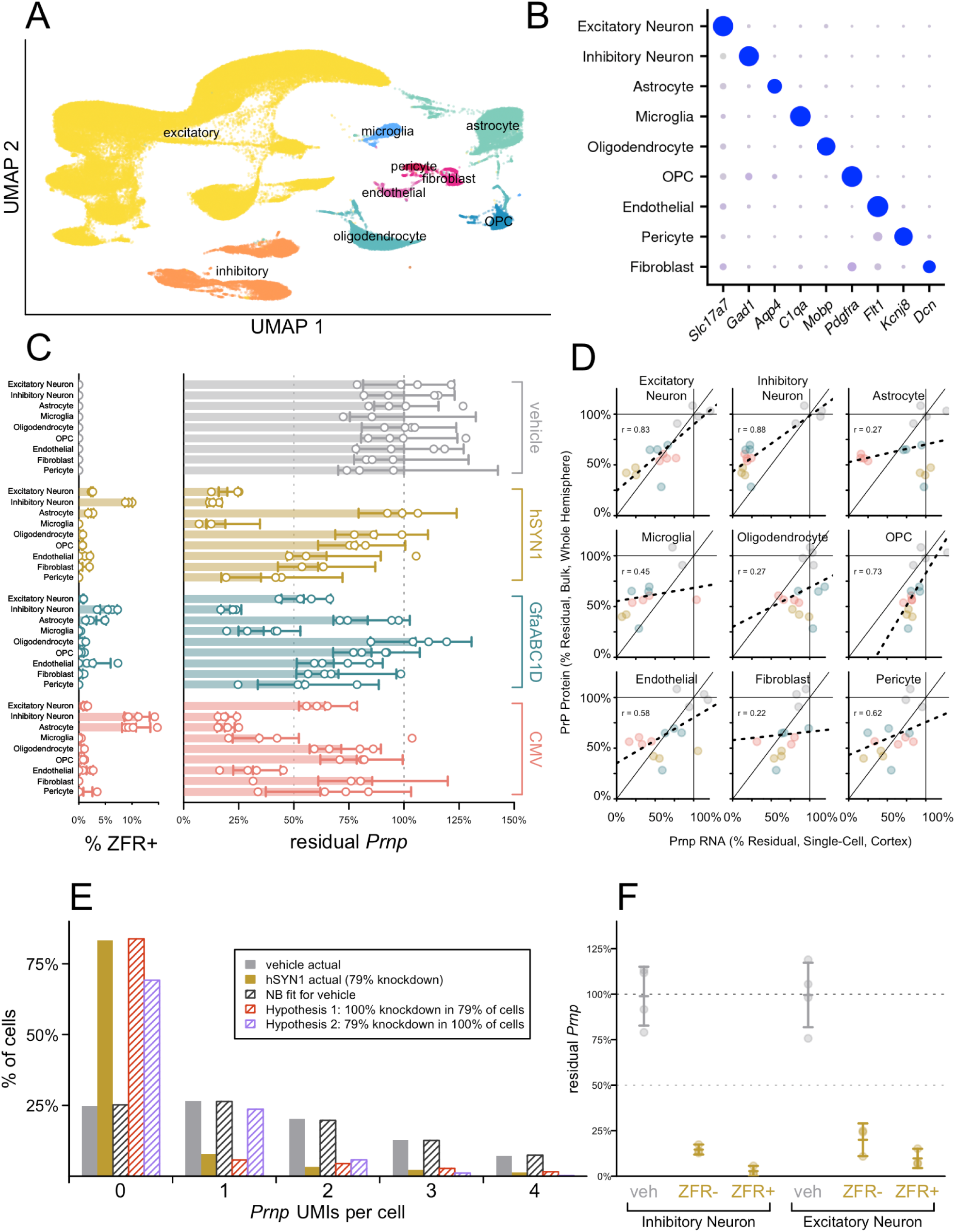
Distribution of mZFR1 and repression of *Prnp* at the single-cell level. **(A)** Two-dimensional Uniform Manifold Approximation and Projection (UMAP) graph shows single-cell gene expression profiles from visual cortex collected for analysis from animals set up in Figure 2A. **(B)** Clustering of selective marker genes used in each assigned cell type are characterized using dot plots. Color gradient represents lower (gray) to higher (blue) expression while dot size represents narrower (small) to broader (large) expression. **(C)** ZFR expression and *Prnp* mRNA expression by cell type and promoter at 5 weeks post-dose. Dots represent individual animals while lines and error bars represent mean and 95% confidence interval (CI) (N = 4 for Vehicle (grey symbols), GfaABC1D (green symbols), and CMV (red symbols), N= 3 for hSYN1 (yellow symbols)). Individual animals with residual ≥150% (apparent artifacts of noise and variability in low N cell types) are plotted at the 150% mark. **(Left panel)** the percentage of cells in which at least one Unique Molecular Identifier (UMI) of ZFR was detected. (**Right panel)** Residual *Prnp* UMIs per million, normalized to the mean of the vehicle group set to 100%. **(D)** Correlation of cell type specific residual *Prnp* mRNA (x-axis) at the single cell level in cortex from Figure 3C and PrP protein expression level (y-axis) of the whole hemisphere at the bulk level from Figure 2C. The best-fit line is shown as the dashed line in each graph with the Pearson’s correlation rho (ρ) value shown above. **(E)** Histogram of *Prnp* UMIs per single neurons (both excitatory and inhibitory) for vehicle (gray) and hSYN1-mZFR1-T2A-GFP-H2B (brown) treated animals. Shaded gray bars indicate the distribution predicted by a negative binomial (NB) model fit to the vehicle data. Shaded red bars indicate the expected NB distribution if the knockdown came from 100% knockdown in a subset of cells; shaded purple bars indicate the expected NB distribution if the knockdown came from equal knockdown in 100% of cells. **(F)** Residual *Prnp* mRNA expression in neurons either from vehicle (veh) or hSYN1-mZFR1-T2A-GFP-H2B treated animals broken down by those in which at least one UMI of ZFR was detected (ZFR+) or not detected (ZFR-). Note that a majority of ZFR-cells were likely transduced by AAV and expressing mZFR1 because a low residual *Prnp* level is observed; however, no detection of UMI for mZFR1 might be due to the assay sensitivity limit from sparse single nucleus sequencing.

The overall pattern of *Prnp* mRNA repression across cell types mirrored the distribution of ZFR+ cells (Figure 3C). *Prnp* knockdown in both excitatory and inhibitory neurons was highest for the treatment group receiving the hSYN1 promoter construct similar to the results of ISH/IHC. The construct with GfaABC1D promoter was surprisingly inefficient in astrocytes. The construct with CMV promoter was efficient in expressing the transgene in both inhibitory neurons and astrocytes. Transgene expression was not detected in microglia, consistent with the known tropism of AAV-PHP.B (Deverman et al. 2016); however, *Prnp* levels were lower in microglia for all three promoters compared to vehicle (Figure 3C). The highest reduction in PrP protein was noted for the hSYN1 promoter construct as measured by ELISA in the whole brain hemisphere (Figure 2C), and when *Prnp* repression for each cell type was compared to bulk PrP reduction, neurons yielded the tightest correlation (rho = 0.83 and 0.88, Pearson’s correlation) (Figure 3D).

### Binomial Distribution Modeling

To better understand whether bulk knockdown of *Prnp* by mZFR1 arises from complete silencing in a subset of neurons (hypothesis 1) or incomplete silencing across all neurons (hypothesis 2), the distribution of *Prnp* counts per individual neuron was assessed. A negative binomial distribution model, which accounts for random variability in detection of *Prnp* transcripts given sparse single cell sequencing (Figure 3E, black diagonal bar) fit the actual vehicle data (Figure 3E, solid gray bar) very tightly. The actual data from the treated animals (Figure 3E, yellow bar) mirrored hypothesis 1 (Figure 3E, red diagonal bar) with 100% knockdown in a subset of neurons, with no *Prnp* repression in the remaining neurons. In particular, the high number of neurons with zero *Prnp* counts and the low number of neurons with exactly 1 *Prnp* count (<10%) could only be explained by such a model. In contrast, a model (hypothesis 2) in which all neurons have an equal amount of knockdown (Figure 3E, purple diagonals bar) would predict fewer neurons with zero counts and far more with 1 count. This suggests that ZFRs yielded profound, almost complete, knockdown in the cells they reach. This is in contrast to knockdown of *Prnp* gene expression with ASOs (Mortberg et al. 2023), where incomplete silencing was observed across all neurons, consistent with hypothesis 2 (Supplementary Figure 3). When neurons were stratified into those having at least one (ZFR+) or zero (ZFR-) ZFR transcript counts, even the ZFR-category showed substantial knockdown (Figure 3F), suggesting again that this category includes both transduced and untransduced cells. In the ZFR+ category, residual *Prnp* is just 3% of vehicle in inhibitory and 10% in excitatory neurons (Figure 3F) in cortex, again consistent with nearly complete silencing in transduced cells. Because both the ISH and single-cell sequencing analyses indicated that vectors using the hSYN1 promoter yielded the deepest knockdown in neurons, the cell type affected in prion disease, we concluded that driving ZFR expression with the hSYN1 promoter has the greatest potential for disease modification in prion-infected mice.

### Single Treatment with AAV-delivered mZFRs Reduces *Prnp* Expression and Extends Survival in a Prion Disease Mouse Model

A pilot study (Survival Study A) was conducted to assess the impact of two constructs hSYN1-mZFR1 and hSYN1-mZFR2 on survival (Figure 4A). RML-inoculated mice received a 1E+14 vg/kg dose at either 60 dpi, when animals have subclinical pathology, or at 122 dpi, around symptom onset. For mice treated at 60 dpi, survival was significantly extended, with median survival times of >449.5 dpi for hSYN1-mZFR1 and 301 dpi for hSYN1-mZFR2, compared with vehicle-treated control animals (median survival of 157 dpi) (Figure 4B). For mice treated at 122 dpi, median survival time increased from 160 dpi for the vehicle-treated group to 378 dpi for hSYN1-mZFR1 and 490 dpi for hSYN1-mZFR2 (Figure 4C). In contrast, animals treated with a control GFP construct showed a median survival time similar to the vehicle control. Altogether, a subset of hSYN1-mZFR-treated animals (2/4 in the 60 dpi hSYN1-mZFR1 group, 1/5 in the 60 dpi hSYN1-mZFR2 group, and 2/5 in the 122 dpi hSYN1-mZFR2 group) survived to the end of study, although all surviving animals showed disease symptoms by 500 dpi. In addition to the lifespan extension seen with the hSYN1-mZFRs, significant delays in both rise of plasma NfL levels and weight loss were observed (Figure 4D-G).

**Figure 4.**
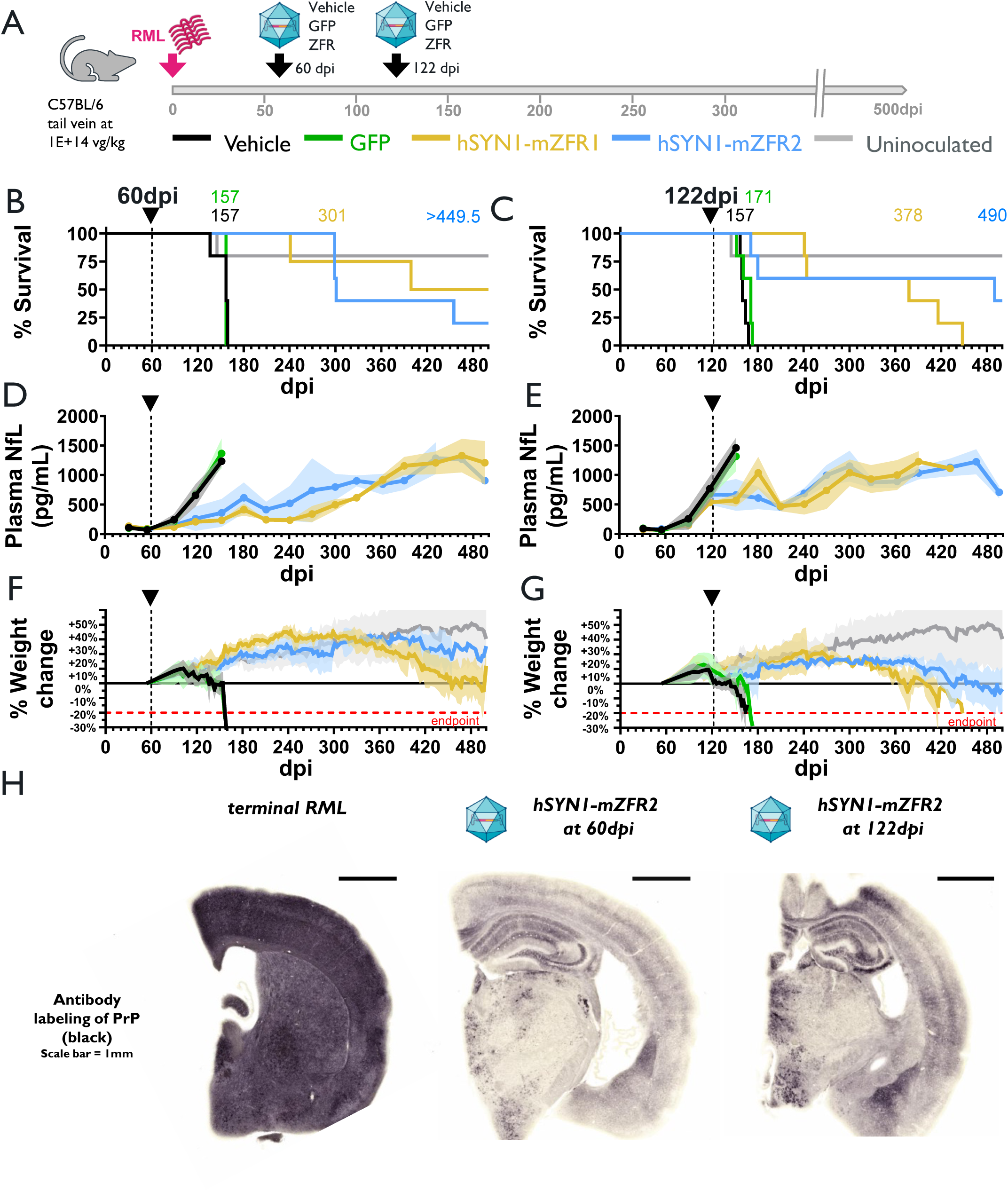
Impact of AAV-hSYN1-mZFR1 treatment on prion disease survival, body weight gain, and plasma NfL levels in RML-inoculated mice. **(A)** Schematic diagram showing experimental design and timeline of Survival Study A. Each cohort is N=4-5 animals, with RML (symbolized as misfolded protein) inoculation on day 0 (pink arrow) through intracerebral injection and treated with 1E+14 vg/kg of AAV-PHP.B-hSYN1-mZFR1 or AAV-PHP.B-hSYN1-mZFR2 via tail vein injection at an early timepoint (60 dpi) when NfL begins to rise **(B, D, F)** or at a late timepoint (122 dpi) near the onset of clinical symptoms **(C, E, G)**. Each timepoint for ZFR, GFP, or vehicle treatment is marked by black arrows. **(B, C)** Survival curves reflecting all-cause mortality. RML-inoculated mice treated with either vehicle, GFP, hSYN1-mZFR1, and hSYN1-mZFR2 are represented by black, green, yellow, and blue lines, respectively. Data from the uninoculated control group is represented by the gray line. In **(C)**, 3 animals were underdosed during the tail vein injection procedure: hSYN1-mZFR1, 2 underdosed animals survived until 378 and 416 dpi, and for hSYN1-mZFR2, 1 underdosed animal survived to 171 dpi. **(D, E)** Plasma NfL concentrations (pg/mL) over the course of the study. Solid lines and shaded areas represent mean and ±SD, respectively. **(F, G)** Percent body weight change relative to individual baseline (body weight at 54 dpi). Solid lines and shaded areas represent mean and ± SD, respectively. **(B-G)** Black downward arrowheads denote time of test articles dosing. **(H)** Representative images with staining for PrP deposition on brain sections from a mouse with terminal prion disease euthanized around 151 dpi, and hSYN1-mZFR2-treated mice from Survival Study A treated at either 60 or 122 dpi, and euthanized at 500 dpi.

We conducted IHC analysis to assess PrP protein aggregates in the brains of ZFR-treated RML-inoculated mice surviving to the scheduled termination date (Figure 4H). An untreated control RML-inoculated mouse euthanized at 151 dpi showed an intense amount of PrP labeling depicting terminal disease stage. In contrast, hSYN1-mZFR2-treated RML-inoculated mice that survived to 500 dpi revealed a profound, though uneven, decrease in PrP staining compared to diseased mice at endpoint. Interestingly, despite showing disease symptoms, the level of PrP protein was not only lower than the pattern seen in the mouse with terminal RML, but also reduced compared to uninoculated, untreated mice at the same age. (Supplementary Figure 4), indicating that ZFR mediated repression persisted for the full study duration.

The dose relationship between PrP reduction and survival benefit was investigated using mZFR1 in Survival Study B. One arm of this study was conducted with hSYN1-mZFR1 administered at doses of 1E+13 (low), 3E+13 (mid), and 1E+14 (high) vg/kg and vehicle (control) 119 days post-RML inoculation (Figure 5A). RML-inoculated mice in all hSYN1-mZFR1-dosed groups showed extended survival with a clear dose-related benefit (Figure 5B). Median survival duration calculated based on all-cause mortality was increased from 169.5 dpi in the vehicle control group to 340 dpi, 462 dpi (2/10 mice survived to end of study), and >494 dpi (median undefined as 5/10 animals survived to end of study) in the low, mid, and high dose groups, respectively. Prior to hSYN1-mZFR1 treatment, all RML-inoculated groups reached an initial plateau in mean body weight gain between 105 to 116 dpi, after which they began to lose weight. Following administration of hSYN1-mZFR1 at 119 dpi, the low dose group recovered to its pre-dosing weight, and the high dose group achieved a maximum weight increase of 19% (Figure 5C). Surviving animals in the low, mid, and high dose groups continued to gain or maintain weight on average through 225, 293, and 405 dpi. When weight loss did eventually occur, its rate was greatly attenuated compared to untreated animals. Increase in all-cause median survival was also observed for RML-inoculated mice treated with hSYN1-mZFR1 on days -21 and 63 dpi; however, the majority of the animals in these groups developed severe dermatitis unrelated to the treatment that required early euthanasia, confounding the survival assessment in these groups (Supplemental Figure 5.1). In addition, dose dependency was observed in general activity and nesting behavior (Supplemental Figure 5.2)

**Figure 5.**
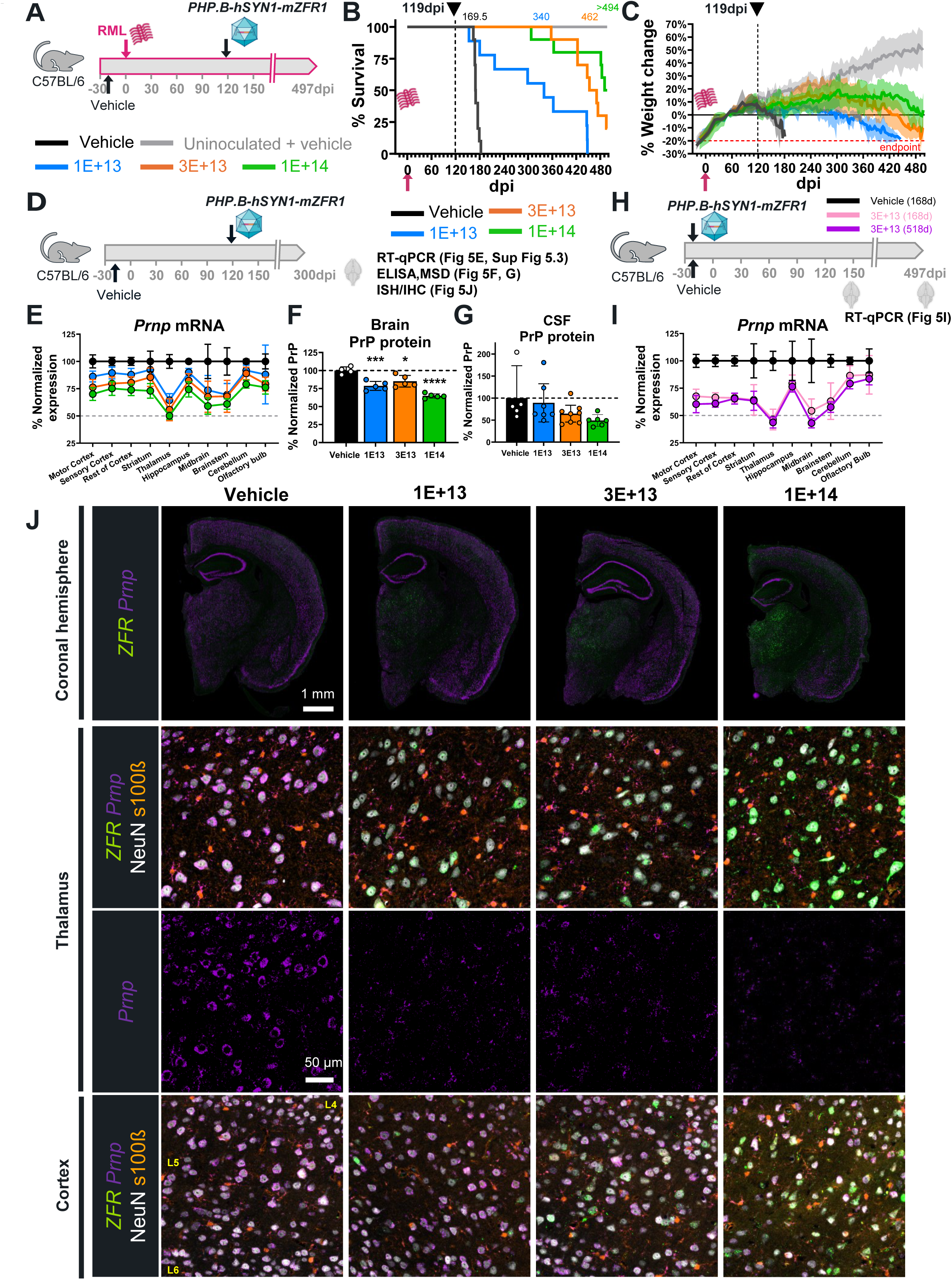
hSYN1-mZFR1 treatment mediates dose-dependent neuronal *Prnp* reduction and survival extension in RML-inoculated mice. **(A)** Schematic diagram showing experimental design for survival monitoring arm in Survival Study B. RML-inoculation day assigned as day 0. Vehicle was dosed at -21 dpi (n=5) and hSYN1-mZFR1 test articles at different dose levels were administered at 119 dpi (n=10/group, except n=9 in low dose 119 dpi group). **(B)** Survival curves representing all-cause mortality. **(C)** Percent body weight change relative to individual baseline at 49 dpi. Solid lines and shaded areas represent means ± SD. Black dashed lines, and the black downward arrowhead denote the dosing of hSYN1-mZFR1 at 119 dpi. **(D)** Schematic diagram showing experimental design for pharmacology analysis arm with necropsy on 301 dpi (denoted by a brain image on the graph) in Survival Study B (n=10 animals/group). Vehicle and hSYN1-mZFR1 test articles were dosed at -21 and 119 dpi, respectively, matching the design used in panel A, except without RML inoculation. **(E)** *Prnp* mRNA expression in different brain regions collected on 301 dpi. The line graph shows mean ± SD values from each treatment group normalized to the vehicle group. Sample numbers: vehicle (n=8), 1E+13 vg/kg (n=10), 3E+13 vg/kg (n=10), and 1E+14 vg/kg (n=10 in all regions except n=9 in brainstem and olfactory bulb). Two-way ANOVA test with Dunnett’s comparison for low-, mid-, and high-dose groups showed significant differences in all brain regions when compared to the vehicle group, except striatum and cerebellum of the low dose group. **(F)** Normalized expression level of PrP protein (mean ± SD) in brain hemisphere. Not all brains collected were used for this analysis; therefore, sample numbers were vehicle (n=4), 1E+13 vg/kg (n=5), 3E+13 vg/kg (n=5), 1E+14 vg/kg (n=5). One-way ANOVA test with Dunnett’s comparison between vehicle and hSYN1-mZFR1-dosed groups. P-values are 0.003 (1E+13 vg/kg), 0.013 (3E+13 vg/kg), and <0.0001 (1E+14 vg/kg). **(G)** Normalized expression level of PrP protein (mean ± SD) in CSF. Sample numbers are vehicle (n=5), 1E+13 vg/kg (n=7), 3E+13 vg/kg (n=(8), 1E+14 vg/kg (n=6) after excluding animals that had insufficient CSF volume collected or had CSF samples with blood contamination. Kruskal-Wallis test with Dunnett’s comparison P-values are >0.99, 0.71, 0.06 for the low, mid, and high dose. **(H)** Schematic diagram showing experimental design for testing durability of hSYN1-mZFR1 in the survival study B. Each cohort was N=10 non-RML inoculated animals. Mouse brains from one group of vehicle-treated (black) and one group of hSYN1-mZFR1-treated (3E+13 vg/kg, pink) were collected 168 days post-dosing. Another group of hSYN1-mZFR1-treated (3E+13 vg/kg, purple) was euthanized for analysis 518 days post-dosing. **(I)** *Prnp* mRNA expression in different brain regions collected as described in H were analyzed by RT-qPCR (mean ± SD) and scaled to the vehicle group mean. Sample numbers were n=10 in most of the brain regions for all three groups except: vehicle (n=9, striatum, brainstem, and cerebellum; n=8 olfactory bulb), 3E+13 vg/kg (168d) (n=9, midbrain; n=8, olfactory bulb), and 3E+13 vg/kg (518d) (n=8, olfactory bulb). Two-way ANOVA test with Dunnett’s comparison was significant for all brain regions compared to the vehicle group. **(J)** Representative microscopy images from brain hemispheres with ISH/IHC analysis for groups demonstrated in Figure 5D. Top row of images: full coronal section demonstrated widespread ZFR (green) expression in brains from hSYN1-mZFR1-treated animals. Second row: overlayed images showing neurons (NeuN, white) in thalamus transduced with hSYN1-mZFR1 (green) expressed less or no *Prnp* mRNA (purple). Astrocytes (s100ß, orange) showed no hSYN1-mZFR1 expression nor *Prnp* reduction comparing hSYN1-mZFR1-treated and vehicle-treated animal. Third row: same region as shown in the second row but showing only the *Prnp* staining. The bottom row shows representative overlayed images from cortex of vehicle and each treatment group. Cortex layer 4, 5, and 6 are labeled as L4, L5, and L6 on the image.

A second arm of Survival Study B was conducted to assess the effect of hSYN1-mZFR1 on *Prnp* mRNA, PrP protein levels, histological endpoints, and survival (Figure 5D-I). Animals not inoculated with RML (uninoculated mice) received hSYN1-mZFR1 at the same dose levels as in the first arm, and treatments were administered to mimic the 119 dpi dosing scheme of the first arm. Brain samples were collected 322 and 182 days after dosing of vehicle or hSYN1-mZFR1, respectively. Dose-dependent reductions in *Prnp* mRNA in various brain regions were observed when compared with the vehicle control group (Figure 5E). For all dose levels tested, markers of neuronal loss and neuroinflammation were comparable to vehicle-treated mice across brain regions (Supplementary Figure 5.3). Dose-dependent decreases in percent normalized PrP protein level in brain hemisphere and CSF were also noted (Figure 5F and 5G, respectively), with the exception that the residual brain PrP level in animals from the mid dose group was slightly higher than in animals of the low dose group. Moreover, *Prnp* mRNA levels in mice treated with hSYN1-mZFR1 after 168 days (Figure 5H, I) were equivalent or slightly higher than those measured after 518 days, indicating that ZFR expression offers stable, long-lasting gene repression in the mouse brain with no diminution of effect for at least 74 weeks. At the single-cell level, ISH/IHC analysis demonstrated an increased number of mZFR1+ neurons with increases in dose. Lower levels of *Prnp* mRNA expression were found in mZFR1+ neurons but not astrocytes in cerebral cortex and deep brain regions like the thalamus (Figure 5J).

### Delivery of a human *PRNP*-targeted ZFR using a novel capsid, STAC-BBB, resulted in widespread transgene expression and *PRNP* mRNA repression in the NHP brain

Having established that ZFR-mediated lowering of *Prnp* can significantly extend survival in RML-infected mice, we investigated whether similar levels of *PRNP* reduction could be achieved in the NHP brain. We developed a highly specific ZFR (hZFR) targeted to a conserved sequence in the human and nonhuman primate *PRNP* gene, which demonstrated >90% repression and high specificity in iPSC-derived human neurons (Supplementary Figure 6). Since the BBB-crossing ability of PHP.B is limited to mice (Huang et al. 2019), we engineered a novel capsid, STAC-BBB, to deliver the hZFR to cynomolgus monkeys. Previously, IV administration of STAC-BBB in a capsid library format resulted in a 700-fold CNS enrichment in neuronal expression compared to its parental serotype, AAV9 (Tiffany et al. 2024). To characterize STAC-BBB tropism, transduction efficiency, and *PRNP* repression in the same animal, we used a ubiquitous CAG promoter to drive expression of a GFP-H2B-T2A-hZFR transgene (Figure 6A). Three NHPs received a 2E+13 vg/kg dose of STAC-BBB CAG-GFP-H2B-T2A-hZFR. After 19 days, brains were collected for IHC, bulk RT-qPCR, and single-cell RNA-scope ISH analyses (Figure 6A). Chromogenic anti-GFP IHC staining on fixed brain tissue sections detected widespread nuclear GFP expression across all cortical and subcortical regions, including the temporal and motor cortices, basal ganglia, thalamus, hippocampus, and brainstem (Figure 6B). Cellular morphology revealed by hematoxylin counterstaining suggested neurons were the primary transduced cell type. Indeed, immunofluorescent staining of GFP and NeuN on the same sections revealed neuronal transduction efficiencies ranging from ∼26-65% in these regions (Figure 6C, D). To evaluate the distribution of STAC-BBB hZFR expression and *PRNP* repression across the rostral-caudal axis, we performed RT-qPCR analysis of ZFR expression levels on 220 tissue punches collected from 35 brain regions spanning 10 brain levels (Figure 6E, top panel). Widespread ZFR transcript expression was detected in all sampled brain levels and regions (Figure 6E, bottom panel). *PRNP* levels were reduced in all 35 brain regions, with relatively consistent performance across animals (Figure 6F). Average bulk *PRNP* transcript reduction ranged from 10-20% in most regions, with the pons, lateral geniculate nucleus, substantia nigra, and thalamus exhibiting the greatest *PRNP* knockdown (Figure 6F).

**Figure 6.**
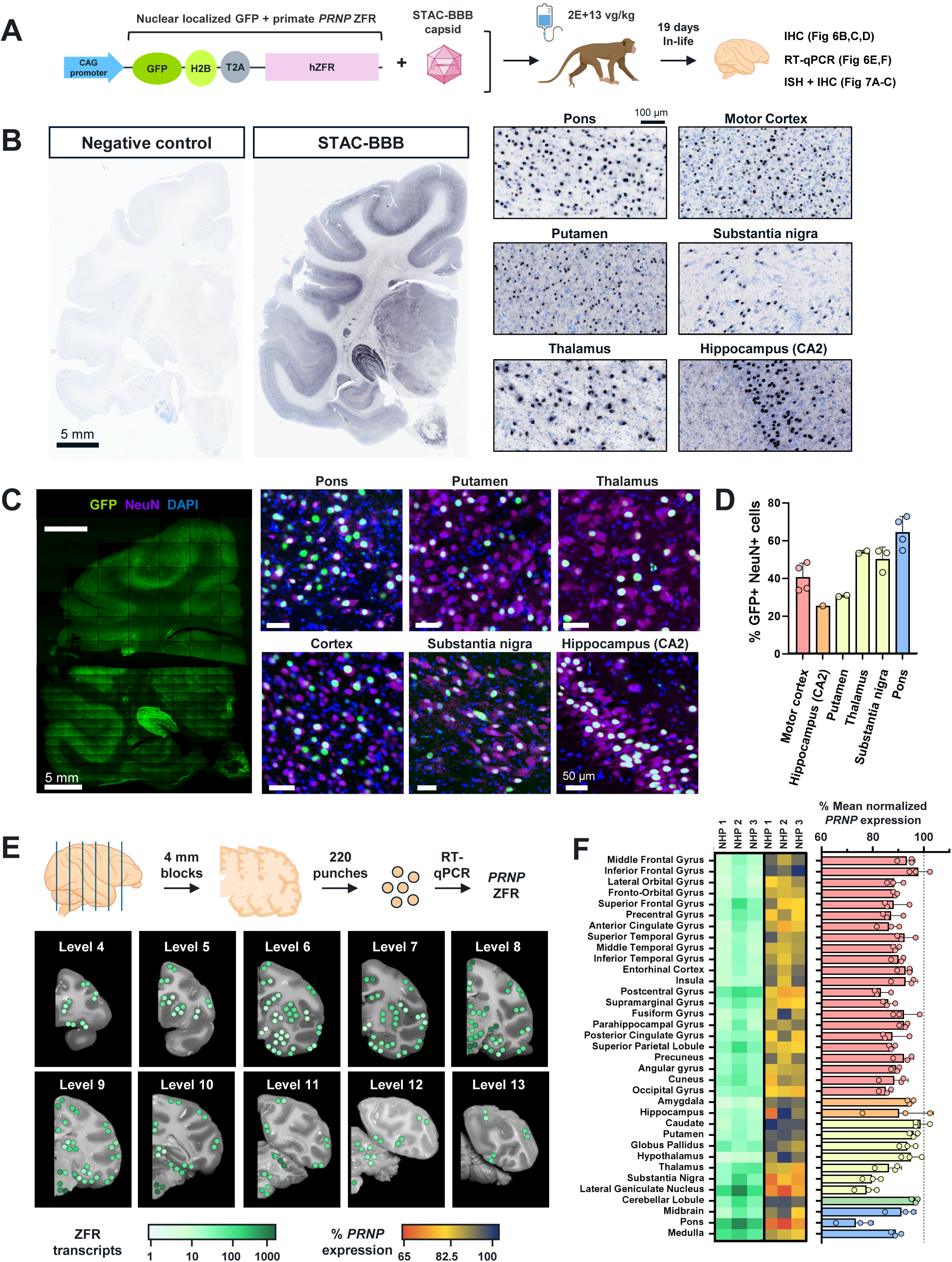
STAC-BBB delivery mediates widespread neuronal transduction throughout the brain in cynomolgus monkeys. **(A)** Schematic diagram of the single-stranded AAV construct that was packaged in STAC-BBB and evaluated in nonhuman primates. (**B)** Representative microscopy images of brain sections stained with an anti-GFP antibody and visualized with chromogenic DAB substrate. The enlarged view highlights the pons, putamen, thalamus, motor cortex, substantia nigra, and hippocampus. Nissl staining (light blue) was used to label all cells. (**C)** Representative multiplexed fluorescence microscopy images of brain sections stained with DAPI (blue), anti-GFP (green), and anti-NeuN (magenta) from STAB-BBB-dosed NHP. Scale bars are 50 µm. (**D)** The percentage of GFP+/NeuN+ cells as quantified in six brain regions. Each dot represents different panels quantified for each region. (**E)** Location of brain punches (circles) analyzed by RT-qPCR overlayed on greyscale images of representative coronal brain sections spanning the rostrocaudal axis. The color of the circle represents the hZFR transcript copy number per ng of total RNA input, with the darker green color indicating higher levels of hZFR expression. (**F)** Heat map (left) demonstrating average expression level of hZFR and *PRNP* mRNA in various brain regions of animals dosed with STAC-BBB-CAG-GFP-H2B-T2A-hZFR as analyzed by RT-qPCR (n=3). Bar graph (right) illustrates the level of *PRNP* expression (mean ± SD) scaled to the average of control-treated animals.

We next characterized hZFR-mediated *PRNP* repression at the single-cell level using dual ISH/IHC. In a vehicle-treated animal, we observed near ubiquitous *PRNP* expression throughout all brain regions examined, with the highest transcript levels detected in NeuN+ cells (Figure 7A, left panels). Representative images from the pons, post central gyrus, midbrain, globus pallidus, and hippocampus show high levels of cytosolic *PRNP* transcripts in neurons. Transcripts associated with neurites and glial cells were also detected, especially in regions with lower neuron density (Fig 7A, left high magnification). In the STAC-BBB hZFR-treated animals, robust neuronal *PRNP* suppression was observed in all regions examined (Figure 7A, right panels). Co-staining for the hZFR transcript and anti-GFP IHC confirmed that a wide range of hZFR expression levels resulted in near-complete removal of *PRNP* from individual neurons (Figure 7A, right high magnification). Quantification of median *PRNP* transcript signal from over 1 million cells from 16 different brain regions revealed an average knockdown of *PRNP* transcripts in NeuN+ cells ranging from 62-96% (Figure 7B). Low or absent *PRNP* signal was only observed in hZFR-treated animals, often in large neurons with minimal or no detectable hZFR or GFP signal (Figure 7A). Indeed, median *PRNP* levels in ZFR-cells exceeded 50% reduction, suggesting that a significant fraction of cells in these regions had near complete reduction of *PRNP* as well. These results are consistent with our observations in mice and indicate that the sensitive ISH and IHC methods used to visualize hZFR expression in NHPs may underestimate the percentage of individual cells that are sufficiently transduced and thereby protected from PrP^Sc^ toxicity.

**Figure 7.**
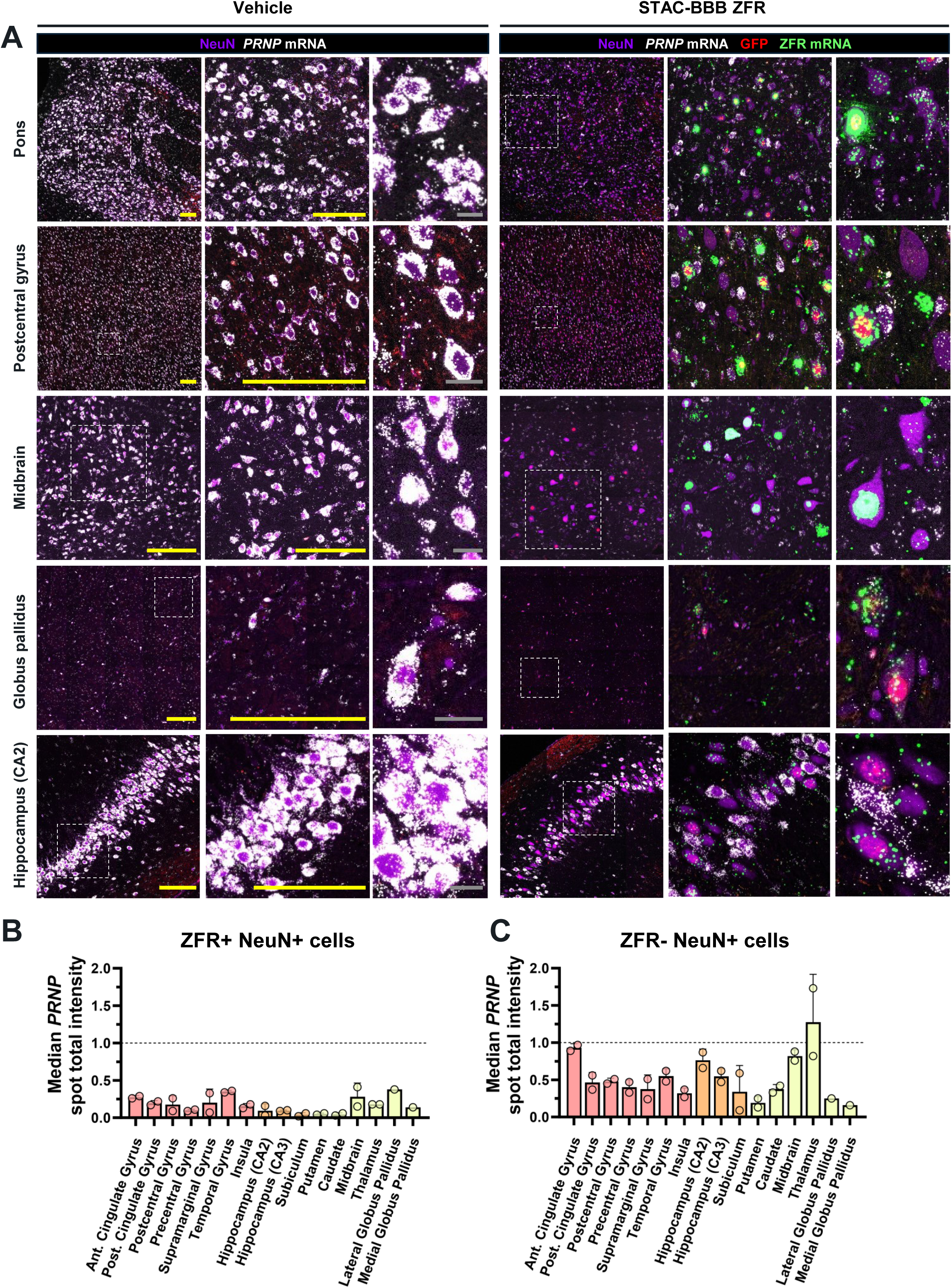
Widespread reduction of *PRNP* mRNA in hZFR-expressing neurons in the nonhuman primate brain. **(A)** Representative microscopy images from brain sections stained for ISH probes: *PRNP* mRNA (white) and hZFR mRNA (green) and IHC antibodies: anti-NeuN (purple) and anti-GFP (red). Images from vehicle-treated (left panels) and hZFR1-treated (right panels) animals for the pons, motor cortex, midbrain, globus pallidus, and hippocampus are shown. Enlarged views of images are shown increasing from left to right. Scale bars in yellow and gray are 200 µm and 20 µm, respectively. **(B, C)** Quantification of *PRNP* mRNA expression at the single-cell level was plotted as median *PRNP* spot total intensity from **(B)** ZFR+/NeuN+ and **(C)** ZFR-/NeuN+ cells from different brain regions (x-axis) of hZFR1-treated (n=2) or control-treated (n=1) animals. Lateral and medial globus pallidus was not present for sections analyzed for one hZFR-treated NHP. Values (mean ± SD) plotted were normalized to values measured from vehicle-treated animals.

## Discussion

In this study, we demonstrate the profound improvement in survival resulting from a one-time IV administered gene therapy in a mouse model of prion disease. While the efficacy of PrP lowering in mouse models of disease has previously been established for a CSF-delivered ASO and divalent siRNA (Minikel et al. 2020; Gentile et al. 2024), our approach has the advantages of ease of administration, constant pharmacologic activity, and broad brain biodistribution achieved via the vasculature, which is expected to scale with brain size. Our data appear to show a greater impact of ZFRs on survival than ASOs despite similar levels of PrP knockdown. For example, ZFR-mediated bulk PrP repression of approximately 35% in whole brain hemisphere was associated with a survival extension of ∼42-47 weeks (Figure 5). In a previous study, transient 33% lowering of *PRNP* transcript in cortex using an ASO delayed a symptomatic endpoint by 6 weeks (Minikel et al. 2020). More uniform suppression across different brain regions and constant silencing (i.e., no peak/trough effects) achieved with ZFRs may be one contributing factor. Another difference is the distribution of activity at the single-cell level. ASOs appear to provide approximately uniform, and incomplete, silencing across all cells within a cell type and brain region (Mortberg et al. 2023); in contrast, ZFR-transduced cells have near complete elimination of *Prnp* transcript, as evidenced by the distribution of *Prnp* counts at the single-cell level and by the near absence (3-10% residual) of *Prnp* in neurons where a ZFR transgene read was detected (Figure 2 and 3). This highlights the importance of AAV distribution throughout the brain, as near-complete repression can be achieved in all transduced cells, consistent with the ZFR having only 2 molecular targets per cell as it acts directly at the DNA level. Given prion neurotoxicity is cell-autonomous and PrP expression-dependent (Brandner, Isenmann, et al. 1996; Brandner, Raeber, et al. 1996; Lakkaraju et al. 2022), cells with completely silenced PrP should be fully resistant to prion disease. This may explain why we observed animals surviving to end of study, even for post-symptomatic ZFR treatments at 119 and 122 dpi, an outcome not previously reported for any other agent.

The timing of treatment was previously identified as a critical factor for survival extension in prion disease models (Minikel et al. 2020), where time to a symptomatic endpoint was markedly improved when ASOs were dosed at -7 dpi to78 dpi, but minimally impacted when dosed at 105 dpi onwards. In contrast, we observed remarkable survival extensions for all AAV-ZFR doses tested regardless of when they were given, even at the latest symptomatic intervention timepoints (119 and 122 dpi) (Figure 4C and 5B). These results demonstrate the potential of AAV-ZFRs for treating all forms of symptomatic prion disease, which accounts for nearly all diagnosed cases, including sporadic, familial, and transmissible forms. For asymptomatic carriers of disease-causing *PRNP* mutations, preventative PrP lowering is expected to be the most impactful; however, the unpredictable age of onset (Minikel et al. 2019) and current lack of definitive prognostic biomarkers (Mok et al. 2023; Vallabh et al. 2024) mean that prophylactic dosing may occur years or even decades before clinical symptoms arise (Vallabh et al. 2020). Therapies requiring lifetime repeat intrathecal injections, for example, biweekly or monthly ASO administration (Miller et al. 2022; Mummery et al. 2023), could be burdensome, particularly for asymptomatic patients. A one-time IV delivery of an AAV-ZFR could provide substantial preventative benefits, as evidenced by the durable effects observed in our studies (lasting at least 74 weeks in mice) (Figure 5I). In summary, our findings support an AAV-ZFR gene therapy as a viable potential treatment for all forms of prion disease in symptomatic patients and presymptomatic carriers.

We sought to address one possible risk of an AAV-based ZFR therapy, which is the potential for loss of transgene expression and eventual diminution of target engagement. In previous studies, optimized ZFRs targeting other genes achieved persistent, stable expression in the mouse brain for at least 48 weeks, and maximal, sustained repression was established as early as 7 days post-dosing (Wegmann et al. 2021; Zeitler et al. 2019). Our current study extends stable ZFR-mediated gene repression to the longest reported timepoint of 74 weeks post-dosing. We observed statistically indistinguishable levels of *Prnp* knockdown and ZFR expression across the brain for animals dosed at 5 weeks of age and analyzed either 24 or 74 weeks later (Figure 5I). Additionally, mice surviving to their scheduled termination at 500 dpi exhibited markedly reduced brain PrP levels compared to untreated controls at 151 dpi (Figure 4H). Together with data from NHPs indicating AAV gene therapy persistence for up to 15 years (Hadaczek et al. 2010; Sehara et al. 2017), our findings support the potential for durable, ZFR-mediated *Prnp* repression in the brain. Of note, two recent studies have sought to establish irreversible inactivation of prion expression in vivo. In the first, permanent epigenome modification via recruitment of DNA methyltransferases (DNMTs) was employed to silence *Prnp* in non-inoculated wildtype mice only (Neumann et al. 2024); however, no evaluation of efficacy, durability of effect beyond 6 weeks, or performance in large animals has been reported. In the second, prophylactic co-administration of two AAV vectors was used to deliver a bacterially-derived base editor to a humanized mouse model, conferring a survival extension of ∼50% (An et al. 2025). Given the relatively limited survival benefit observed in a preventative dosing paradigm, requirement for high doses of two AAV vectors, and risk of irreversible off-target effects caused by lifelong editor expression in the brain, more work may be needed to understand the translational potential of this approach.

Whereas ASOs have relatively broad distribution across cell types within a brain region (Mortberg et al. 2023), our study is the first to directly investigate neuron-restricted PrP lowering using a potentially therapeutic modality. *In vivo*, prions can replicate in both neurons and astrocytes (Aguzzi and Liu 2017; Lakkaraju et al. 2022), but not in microglia or oligodendrocytes (Priller et al. 2006; Prinz et al. 2004). Prion neurotoxicity is cell autonomous and requires neuronal PrP (Brandner, Isenmann, et al. 1996; Lakkaraju et al. 2022), and neuron-specific knockout is sufficient to halt prion disease (Mallucci et al. 2003). When *Prnp* was floxed in a Cre-NFH model, PrP protein concentration in the brain appeared to drop by ∼90% (Mallucci et al. 2003), suggesting that even though *Prnp* mRNA is expressed ubiquitously across various brain cell types, PrP protein is produced predominantly in neurons. We found that PrP knockdown in whole brain hemispheres was most tightly correlated with neuronal *Prnp* mRNA knockdown (Figure 2 and 3), consistent with brain PrP being predominantly neuronal in origin, even though the *Prnp* transcript is expressed in all cell types. Therefore, we hypothesized that neuronal PrP reduction would be most critical for prion disease. Our data show that a neuron specific (hSYN1) promoter-driven mZFR specifically silenced *Prnp* in neurons, with limited activity in other cell types (Figure 2 and 3), led to a significant increase in median survival, with some animals surviving to the scheduled end of both survival studies (Figure 4 and 5). These results reinforce existing findings from animal models that suggest lowering *Prnp* expression in neurons is sufficient to slow or prevent disease progression. In addition, use of a cell-type specific promoter combined with the neurotropic, liver-detargeted properties of STAC-BBB in NHP (Tiffany et al. 2024) reduces the risk of possible toxicity due to transgene expression in peripheral tissue, as has been associated with some AAV-based therapies (Hordeaux et al. 2020; Van Alstyne et al. 2021).

Because a substantial fraction of *Prnp* transcripts is produced in glia and other non-neuronal cells, we used multiple approaches to assess knockdown within individual neurons. Data from our mouse studies revealed that *Prnp* transcript levels are reduced by 90-97% in transduced neurons, whereas bulk transcript levels are reduced by ∼10-70% depending on the brain region, AAV dose, and promoter. In NHPs, *PRNP* expression in most cortical regions was reduced by only 10-20% in bulk cell analyses, yet at the single-cell level, the median knockdown effect was ∼75-90% in ZFR+ neurons; in neurons classified as ZFR-, we observed a median knockdown of ∼50% in most cortical regions compared to control levels. This suggests that our ability to detect ZFR expression underestimates the proportion of cells expressing sufficient levels of ZFR to repress *PRNP*. Collectively, these findings support the notion that ZFRs act in a digital fashion at the single-cell level and highlight transduction efficiency as a critical determinant of the number of cells protected from toxic PrP^Sc^.

The unprecedented survival benefit conferred by mZFRs in the RML disease model motivated the development of an hZFR specifically targeting the human and NHP *PRNP* gene (Supplementary Figure 6.1) for evaluation in NHPs. Numerous studies have shown that high-dose AAV administration via intra-CSF routes yield sparse and primarily perivascular transduction of the cortex (Bey et al. 2020; Meseck et al. 2022; Bharucha-Goebel et al. 2024). No CSF dosing route approaches the degree of parenchymal access afforded by the brain capillary network, particularly for more challenging subcortical regions (Pardridge 2016). Additionally, the complexity of certain CSF AAV administration procedures, such as intracisternal injections, could pose unique challenges for CJD patients and would require qualified neurosurgical centers for delivery. To overcome these limitations, we utilized STAC-BBB, a novel AAV capsid that can efficiently cross the BBB in NHPs (Tiffany et al. 2024). STAC-BBB enabled efficient delivery of the hZFR to all 35 distinct brain regions evaluated, resulting in *PRNP* repression at both bulk and single-cell levels (Figure 6 and 7). To our knowledge, this represents the first successful delivery of an epigenetic regulator to NHPs via an IV-administered, BBB-crossing capsid. Potent *PRNP* reduction was observed in transduced neurons across key brain regions, including the cerebral cortex, basal ganglia, thalamus, brainstem and hippocampus – areas where vacuolation, spongiform degeneration, and PrP^Sc^ pathology are commonly observed in postmortem CJD brains. Notably, STAC-BBB-mediated hZFR biodistribution, transduction and pharmacology in NHPs closely mirrored measurements of the same parameters for mZFRs delivered by PHP.B to wildtype and RML-inoculated mice. The similar performance across capsids and ZFRs provides a framework for approximating the level of bulk *PRNP* repression necessary to potentially achieve therapeutic benefit for a STAC-BBB-delivered hZFR. Importantly, our study used a substantially lower IV dose in NHPs (2E+13 vg/kg) than approved AAV gene therapies for human (1E+14 vg/kg for onasemnogene abeparvovec). Higher doses could further boost neuronal transduction and increase neuronal PrP^C^ knockdown. In addition, further engineering of the STAC-BBB capsid could lead to enhanced CNS delivery at similar or reduced doses.

A potential limitation of this work is the use of surrogate mZFR reagents to evaluate pharmacology and efficacy in RML-inoculated mice. We initially planned to use humanized *PRNP* transgenic mice infected with human prions to directly assess the efficacy of hZFRs. However, we abandoned this approach following the documented deaths of laboratory personnel following occupational exposure to human PrP^Sc^ (Brandel et al. 2020; Casassus 2021). Currently, no alternative disease models are available that allow hZFR testing with a prion strain that does not pose an unacceptable risk to human health. To address this limitation, we leveraged the conserved mechanism of action and comparable on- and off-target profiles of the mZFRs and hZFR, and designed our efficacy studies to enable bridging across common measures of transduction, ZFR expression and *PRNP* repression at the bulk and single-cell levels.

Collectively, our results provide strong support for the further development and clinical testing of an IV-administered AAV-ZFR medicine to potentially treat all forms of human prion disease.

## Methods

### ZFR and STAC-BBB capsid sequences

mZFR1, mZFR2, mZFR3, hZFR and STAC-BBB – Amino acid sequences to be provided upon peer-reviewed journal publication.

### In Vitro Experiments

#### Mouse N2a neuroblastoma cell nucleofection

Mouse N2a neuroblastoma cells (ATCC, CCL-131) were cultured according to the manufacturer protocol. For nucleofection, 100,000 N2a cells per well plated on 96-well plates and were resuspended in Amaxa® SF solution. The cells were then mixed with ZF-Rs mRNA at 3 different doses: 100, 300, 1000 ng per well, and transferred to Amaxa® shuttle plate wells. Then the cells were transferred to a 96-well tissue culture plate and incubated at 37°C for 20 hours. Cells were harvested 24 hours post-transfection and processed for RT-qPCR analysis.

#### In vitro transduction of AAV6-ZFRs in mouse cortical neurons

For target gene expression analysis, mouse primary cortical neurons (MCN) (Gibco, A15586) were seeded at 4.00E4 per well (96-well plate) and transduced with AAV6-ZFRs at the following MOIs: 1.00E+2, 3.00E+2, 1.00E+3, 3.00E+3, 1.00E+4, and 3.00E+4 vector genomes (vg)/cell 24 hours after plating. Cells were harvested 7 days after AAV viral transduction, total RNA was isolated, and RT-qPCR was performed. For differential gene expression (microarray) analysis, the cells were seeded at 2.00E5 per well (24-well plate) and transfected with 3.00E+3 vg/cell 48 hours after plating and harvested 7 days after viral transfection.

#### *In vitro* transduction of AAV6-ZFRs in human iPSC-derived GABAergic neurons

For on-target analysis, human iPSC-derived GABAergic neurons (iGABA) (Cellular Dynamics International, R1013) were seeded at a density of 4.00E+4 cells per well of a 96-well plate coated with Poly-L-Ornithine (PLO)/Laminin (Corning, #354657). Cells were transduced with AAV6-ZFR at MOIs from 1.00E+3 to 3.00E+5 vg/cell 48 hours post-plating. For differential gene expression (microarray) analysis, 2.50E5 cells per well were seeded on a PLO/Laminin-coated 24-well plate (Corning, #354659) for 48 hours then transduced with AAV6-ZFR at 1.00E+5 vg/cell. Cells were harvested 19 days after AAV transduction.

### Animal Studies

#### Ethical Statement for Animal Studies

One mouse promoter comparison study was conducted at Atuka Inc. (Toronto, ON, Canada), one mouse pharmacology and two mouse survival studies were conducted at the Broad Institute (Boston, MA, USA), and one nonhuman primate study was conducted at Charles River Laboratories (CRL) Montreal ULC (Senneville, QC, Canada). All facilities were accredited by Association for Assessment and Accreditation of Laboratory Animal Care (AAALAC). All protocols were approved by the respective Institutional Animal Care and Use Committees (IACUC). Studies were performed either in conformance with the U.S. Public Health Policy on the Care and Use of Animals as defined in the Guide to the Care and Use of Animals or Canadian Council on Animal Care (CCAC) guidelines. Species specific standard procedures and conditions for animal care, housing, access to water and food, environment, and room maintenance were used. All other procedures were performed in accordance with laboratory standard operating procedures and/or established laboratory best practices.

#### Pharmacology Study in Mice

A 5-week pharmacology study was conducted in female C57BL/6 mice at the Broad Institute (Figure 1D, E). C57BL/6J mice were purchased from Jackson Laboratory at 5 weeks of age. Mice were intravenously (IV) dosed on Day 0 with either vehicle (formulation buffer, PBS + 0.001% Pluronic F-68, pH 7.1; n=8), AAV-PHP.B.hSYN1.GFP (n=6), AAV-PHP.B-hSYN1-mZFR1 (hSYN1-mZFR1; n=5), or AAV-PHP.B-hSYN1-mZFR2 (hSYN1-mZFR2; n=4) at a dose of 1E+14 vg/kg. On Day 35 post-dose administration animals were euthanized by perfusion with CO2 and brain tissues were collected for analysis. The right brain hemispheres were treated with RNAlater and individual regions were dissected and frozen for molecular analysis (RT-qPCR). Left brain hemispheres from vehicle and ZFR groups only were frozen for PrP protein assessment by ELISA.

#### Promoter Comparison Study in Mice

A 5-week promoter comparison study was performed in wildtype C57BL/6 mice at Atuka Inc. (Figure 2A). C57BL/6NCrl mice (n=4/sex/group) were purchased from CRL at 8-10 weeks of age. Mice were treated IV on Day 1 with a dose of 1E+14 vg/kg of either AAV-PHP.B-hSYN1-mZFR1 (hSYN1-mZFR1), AAV-PHP.B-GfaABC1D-mZFR1 (GfaABC1-mZFR1), or AAV-PHP.B-CMV-mZFR1 (CMV-mZFR1). The control group received vehicle (1X DPBS (+Ca/+Mg) + 0.001% Pluronic F-68, pH 7.2). Following collection of CSF on Day 36, mice were transcardially perfused with 0.9% RNAse–free saline and CNS tissues (brain and spinal cord) and peripheral tissues (liver, heart, spleen, lung, kidney, muscle, and gonads) were collected. The right brain hemispheres of all animals were treated with RNAlater, and individual regions were dissected and frozen for molecular analysis (RT-qPCR). Left brain hemispheres from 2 animals/sex/group were frozen and used for PrP protein analysis by ELISA and single nucleus RNA-seq (snRNA-seq) analysis. The left brain hemispheres from the remaining animals in each group were preserved in 10% neutralized buffered formalin (NBF) at room temperature for 24 hours, then transferred to 70% ethanol and embedded in paraffin within 7 days following transfer to ethanol, for *in situ* hybridization / immunohistochemistry (ISH/IHC; RNAscope) evaluation.

### Mouse Survival Studies

Two survival studies (Survival Study A and B) were conducted at the Broad Institute.

#### Intracerebral Inoculation with RML Prions (Prion Challenge)

For both survival studies C57BL/6N mice were infected by intracerebral prion inoculation with 30 μL of a 1% brain homogenate as previously described [cite PMID: 31361599, 32776089]. Briefly: brains were homogenized (10% wt/vol) in PBS (Gibco, 14190) by 3 x 40s high pulses in 7 mL tubes with zirconium oxide beads (Precellys, KT039611307.7) in a Minilys homogenizer (Bertin Technologies EQ06404-200-RD000.0), then diluted to 1% in more phosphate-buffered saline (Gibco, 14190), irradiated with 7.0 kGy of X-rays on dry ice, and extruded through progressively finer blunt needles (Sai Infusion, B18, B21, B24, B27, B30). Homogenates were then pipetted into glass vials and injected through 31G disposable syringes (Becton Dickinson, 328449) into sealed amber glass vials. Inoculation was freehand between the right ear and midline, under isoflurane anesthesia.

#### Survival Study A

In Survival Study A (Figure 4A), 8-week-old female C57BL/6N-Elite mice (CRL; n=5/group) were inoculated with RML prion on study Day 0. The mice then received a single tail-vein injection of either vehicle (control group), or AAV.PHP.B.hSYN1.GFP (AAV-hSYN1-GFP), AAV-hSYN1-mZFR1, or AAV-hSYN1-mZFR2 at a dose of 1E+14 vg/kg on 60 or 122 days post inoculation (dpi). Mice were observed twice daily for mortality/morbidity (survival) for up to 500 days and the following parameters were evaluated: median survival, body weights, body weight changes, nesting behavior, and activity. Blood was also collected monthly and processed to plasma for assessment of neurofilament light chain (NfL) levels. Animals were euthanized by CO_2_ inhalation when they met pre-defined endpoint criteria or at study termination (500 dpi). No tissues were collected from mice euthanized for prion endpoint. For immunohistochemistry (IHC) analysis presented in Figure 4H and Supplementary Figure 4, RML-mice with ZFR treatment and uninoculated, untreated mice that survived until the end of the study (500 dpi) were perfused and brain hemispheres were collected. Additional mouse brain hemispheres were collected from a wildtype C57BL/6 mouse inoculated with RML and euthanized on 151 dpi. Brain hemispheres collected were post-fixed in formalin, followed by treatment with ≥95% formic acid to inactivate PrPSc, transferred to formalin, and then stored in PBS + 0.02% sodium azide, prior to embedding in gelatin blocks. Gelatin embedding and IHC with anti-PrP EP1802Y (Abcam, #52604) were performed at NeuroScience Associates (NSA, Knoxville, TN, USA).

#### Survival Study B

Survival Study B involved a one-time retro-orbital administration of either vehicle control or hSYN1-mZFR1 at different dose levels. In Survival Study A, tail-vein injections were identified as a potential factor of dosing variability; we therefore switched to retro-orbital injections after demonstrating improved dosing consistency in a small pilot study. Female C57BL/6N-Elite mice (CRL) were 7-11 weeks of age at day of RML inoculation (Day 0). The study duration was 518 days.

In one arm of the study, group designations were as follows: one group (5 mice) received vehicle on -21 dpi but were not inoculated. A second group of 10 mice received vehicle on -21 dpi and were inoculated with RML on Day 0. Another group of 10 mice received test article (3E+13 vg/kg) on -21 dpi and were inoculated with RML on Day 0. On 63 and 119 dpi, three groups of 10 mice per dpi day who were inoculated with RML on Day 0 then received 1E+13, 3E+13, and 1E+14 vg/kg of the test article. All animals were monitored for mortality/morbidity (survival) throughout the study for up to 518 days. The same parameters as in Survival Study A were assessed, except NfL levels were not evaluated in this study. Animals were euthanized by CO_2_ inhalation when they met pre-defined endpoint criteria or at study termination (497 dpi).

A second arm in this study was dedicated to assessment of mZFR, *Prnp* mRNA expression, and PrP protein levels. In this arm, animals were not inoculated with RML. Four groups of C57BL/6N mice (n=10/group) received either vehicle control on Day -21 (relative to Day 0), or 1E+13, 3E+13, or 1E+14 vg/kg of the test article on Day 119 (relative to Day 0). The in-life duration for this arm was 301 days post Day 0. One additional group of 10 mice received test article at 3E+13 vg/kg on Day -21 and was allowed to survive until Day 497 (relative to Day 0) to match the study duration in the first arm. On Day 301 or 497 (relative to Day 0), CSF was collected for PrP measurements prior to euthanasia by total body perfusion using PBS. Following euthanasia brain was collected and processed similarly as described for the promoter comparison study discussed above, with right brain hemispheres from all mice designated for molecular (RT-qPCR) analysis, left brain hemispheres from 5 mice/group designated for protein analysis (ELISA), and left brain hemispheres from 5 mice/group evaluated using ISH/IHC.

#### Animal Monitoring

The monitoring scheme for prion-infected mice was the same as described previously (Minikel et al. 2020). Mice were monitored daily for general health. Nest-building behavior, reduced activity, and body weights were monitored at 54 dpi, then weekly from 89 – 115 dpi, and thrice weekly starting at 120 dpi in Survival Study A. In Survival Study B, nest-building behavior and reduced activity were monitored weekly starting from 90 dpi, escalating to thrice weekly at 120 dpi; body weights were monitored weekly starting at -14 dpi, and thrice weekly starting at 120 dpi. Nest-building behavior for each cage was rated for both cotton square nestlets (Ancare) and Enviro-dri packed paper (Shepherd) on a scale of 0 to 2: 0= unused; 1= used/pulled apart, but flat; 2= pulled into a three-dimensional structure. Intermediate scores of 0.5 and 1.5 were permitted. Nestlet and paper scores were averaged to yield a combined score. Submandibular vein bleeds were obtained monthly into lithium heparin vials (BD 365985), kept on wet ice and then spun at 1,000 x G for 15 minutes to isolate plasma for NfL analysis (Survival Study A only). NfL protein was quantified using Simple Plex Human NF-L assay kit on an Ella Automated Immunoassay System (Bio-Techne) according to manufacturer instructions. Animals were euthanized using CO_2_ at 20% weight loss relative to their baseline (taken at 16 weeks of age, corresponding to 54 dpi in Survival Study A and to 49 dpi in Survival Study B) or if moribund (meaning unable to reach food or water), with a pre-specified takedown at 500 or 497 dpi for surviving animals in Survival Study A and B, respectively. Survival curves reflect all-cause mortality including animals euthanized due to intercurrent severe dermatitis; a second set of survival curves was generated showing mortality with animals that were euthanized for severe dermatitis censored.

### Distribution and Pharmacology Study in Nonhuman Primates

A 19-day AAV vector distribution and pharmacology study was conducted in naïve cynomolgus monkeys (*Macaca fascicularis*; Cambodian origin; CRL, QC, Canada) that were 2.5 years old at time of dosing. Two males and one female received a single intravenous administration of an STAC-BBB GFP-H2B-T2A-hZFR vector encoding a ZFR targeting the human/nonhuman primate prion gene (*PRNP*) at a dose level of 2E+13 vg/kg on Day 1. The construct also encoded a nuclear localized green fluorescent protein sequence under the control of the ubiquitous CAG promoter that allowed detection of the construct at the single cell level. Animals also received daily methylprednisolone sodium succinate at a dose level of 1 mg/kg via intramuscular injection as immunosuppressant for the duration of the study. On Day 19, animals were euthanized by isoflurane anesthesia followed by whole-body perfusion using ice-cold RNAse-free PBS. Brains were divided into 4 mm slices with right hemispheres designated for molecular (RT-qPCR) analysis and left hemispheres for GFP IHC and ZFR/PRNP/NeuN (ISH/ISH/IHC RNAscope) evaluation. Right hemisphere slices were treated with RNAlater, followed by collection of biopsy punches throughout the brain. Left hemisphere slices were preserved in 10% NBF for 24 hours, transferred to 70% ethanol, embedded in gelatin or paraffin and processed to blocks.

### Molecular Analyses

#### Affymetrix analysis for the evaluation of off-targets

To evaluate the off-target impact of the AAV6-ZFRs on global gene expression, we performed microarray experiments on total RNA isolated from MCN and iGABA transduced with AAV6-ZFRs using Clariom™ S Assay HT, mouse (Thermofisher, #902972) and Clariom™ S Pico Assay HT, human (Thermofisher, #902964), respectively. For each replicate 70 ng of total RNA was processed according to the manufacturer’s protocol for sample preparation, hybridization, fluidics, and scanning (Affymetrix MTA1.0 GeneChip arrays, Affymetrix). Fold change analysis was performed using Transcriptome Analysis Console 4.0 (Affymetrix) software. AAV6-ZFR treated samples were compared to samples treated with a non–*Prnp*-targeted AAV6-ZFR. Change in gene expression is reported for transcripts (probe sets) with a >2-fold difference in mean signal relative to control and with the FDR-corrected p-values set to <0.05.

#### RT-qPCR for In Vitro Experiments

After each time point for each cell line, cells were harvested and lysed and reverse transcription was performed using a Cells2CT Custom kit (Thermo Fisher Scientific, 4402955C001) following the manufacturer’s instructions. TaqMan quantitative polymerase chain reaction (qPCR) was used to measure the expression levels of *Prnp*, which were normalized to the geometric mean of the expression levels of the housekeeping genes *Atp5b* and *Eif4a2*. RT-qPCR was performed using Bio-Rad CFX384 thermal cyclers. cDNA was diluted 10-fold in nuclease-free water, and 4 μL of diluted cDNA were added to each 10 μL PCR reaction. Plates were prepared using Tecan automated liquid handling robotics. Each sample was assayed in technical quadruplicate. Custom Taqman primer:probe assays were used in this study (Table 1 for MCN, Table 2 for iGABA neurons). The qPCR cycling conditions were as follows: Qiagen Fast Multiplex master mix → 95°C for 5 min, 95°C for 45 s, 60°C for 45 s, plate read, 40 cycles.

**Table 1:**
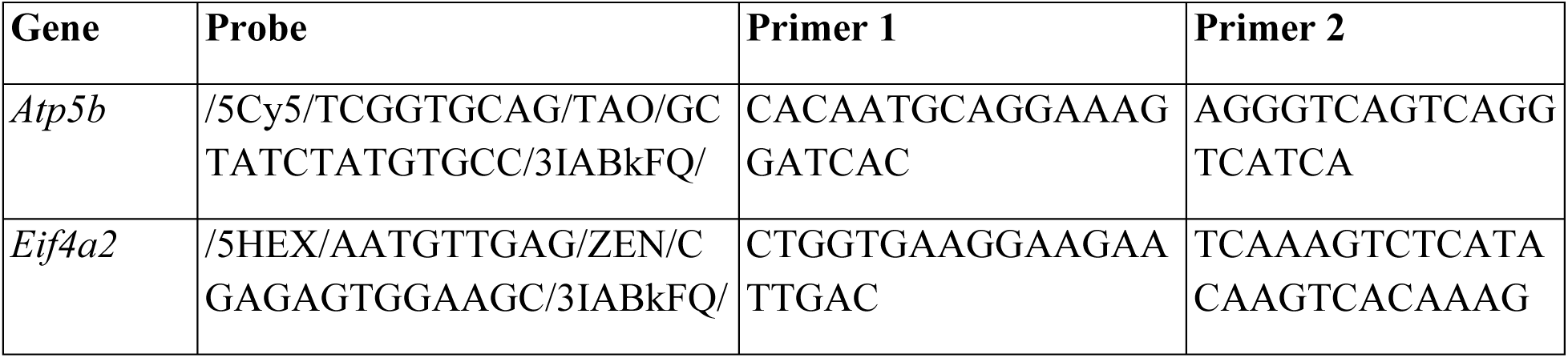
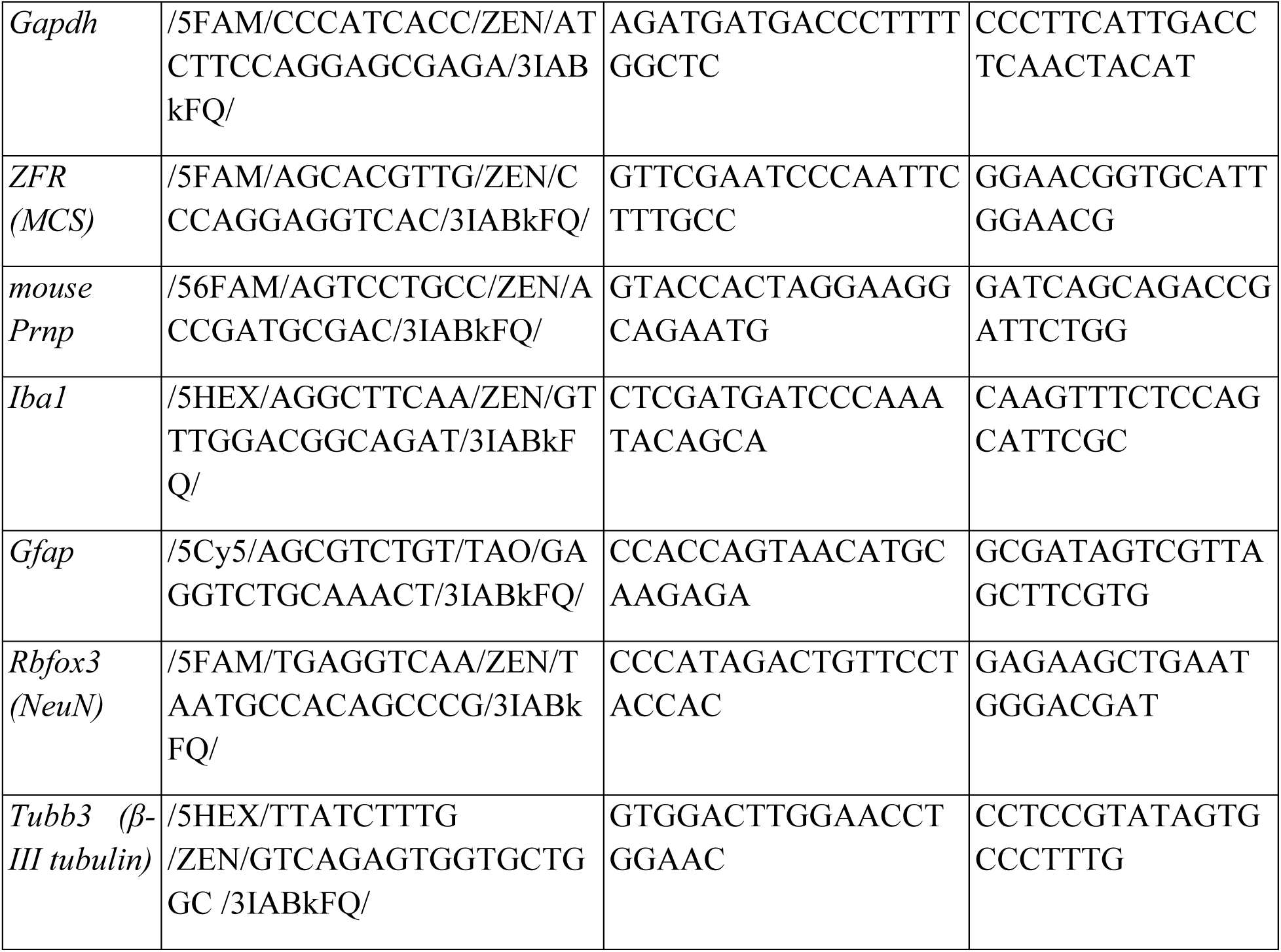
Taqman RT-qPCR Probes (Mouse)

**Table 2:**
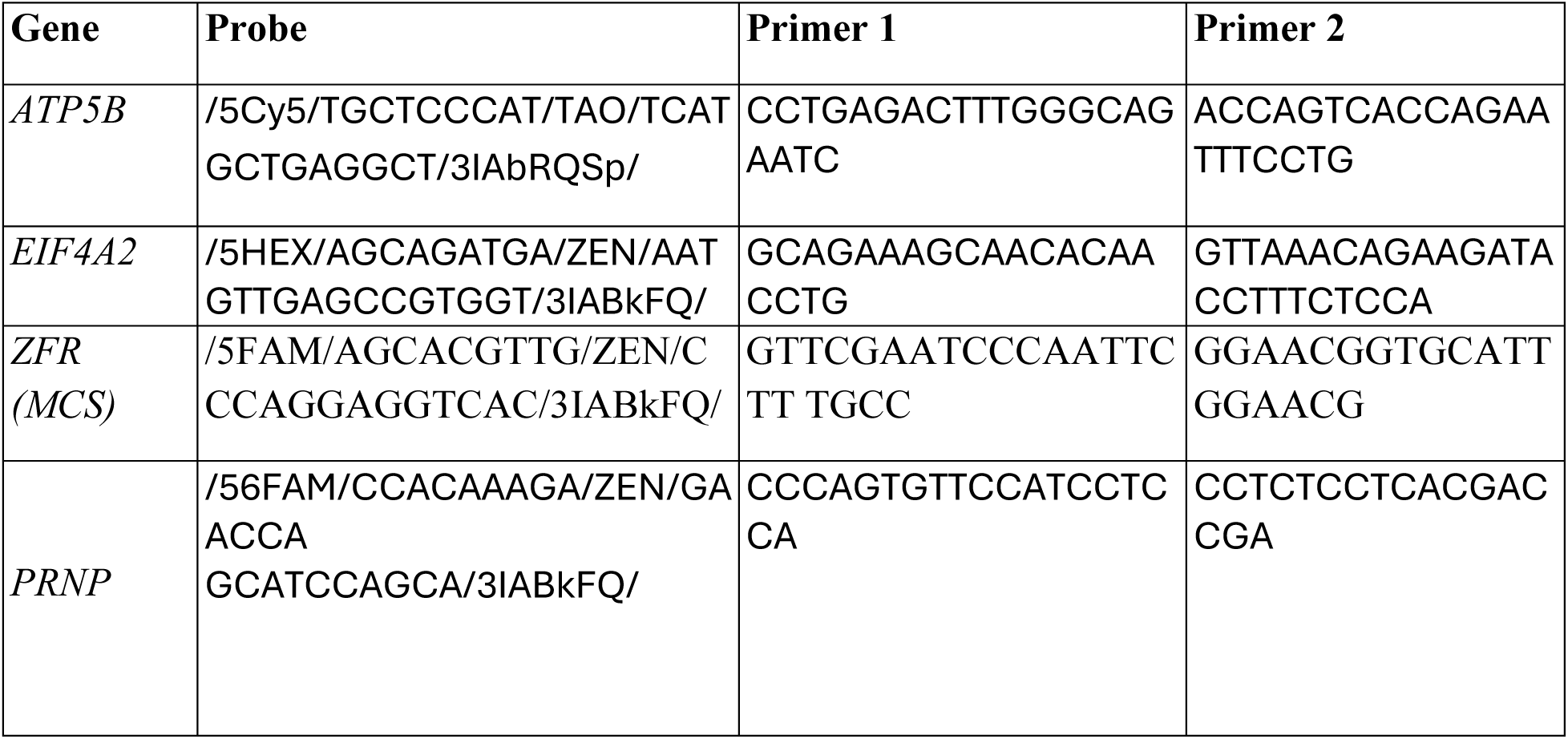
Taqman RT-qPCR Probes (iGABA neuron)

### RNA isolation and RT-qPCR for Animal Studies

RNAlater-preserved brain or peripheral tissues were lysed using a Qiagen TissueLyser and RNA were isolated with MagMax Kit (THermo Fisher, Cat# AM1830). cDNA was prepared using the High-Capacity cDNA Reverse Transcription Kit (Applied Biosystems) using a fixed volume for total RNA input. RT-qPCR was performed using Biorad CFX384 thermal cyclers. cDNA was diluted 20-fold in nuclease-free water, and 4 μL of diluted cDNA was added to each 10 μL PCR reaction. Each sample was assayed in technical quadruplicate. For mouse and NHP studies custom Taqman primer:probe assays were used; the primer/probes sequences are provided in Table 1 and Table 3, respectively, and were ordered from IDT. For both mouse and NHP studies, the RT-qPCR cycling conditions were as follows: Qiagen Fast Multiplex master mix - 95°C for 5 min, 95°C for 45 s, 60°C for 45 s, plate read, 40 cycles; Biorad SsoAdvanced master mix - 95°C for 90 s, 95°C for 12 s, 60°C for 40 s, plate read, 42 cycles.

**Table 3:**
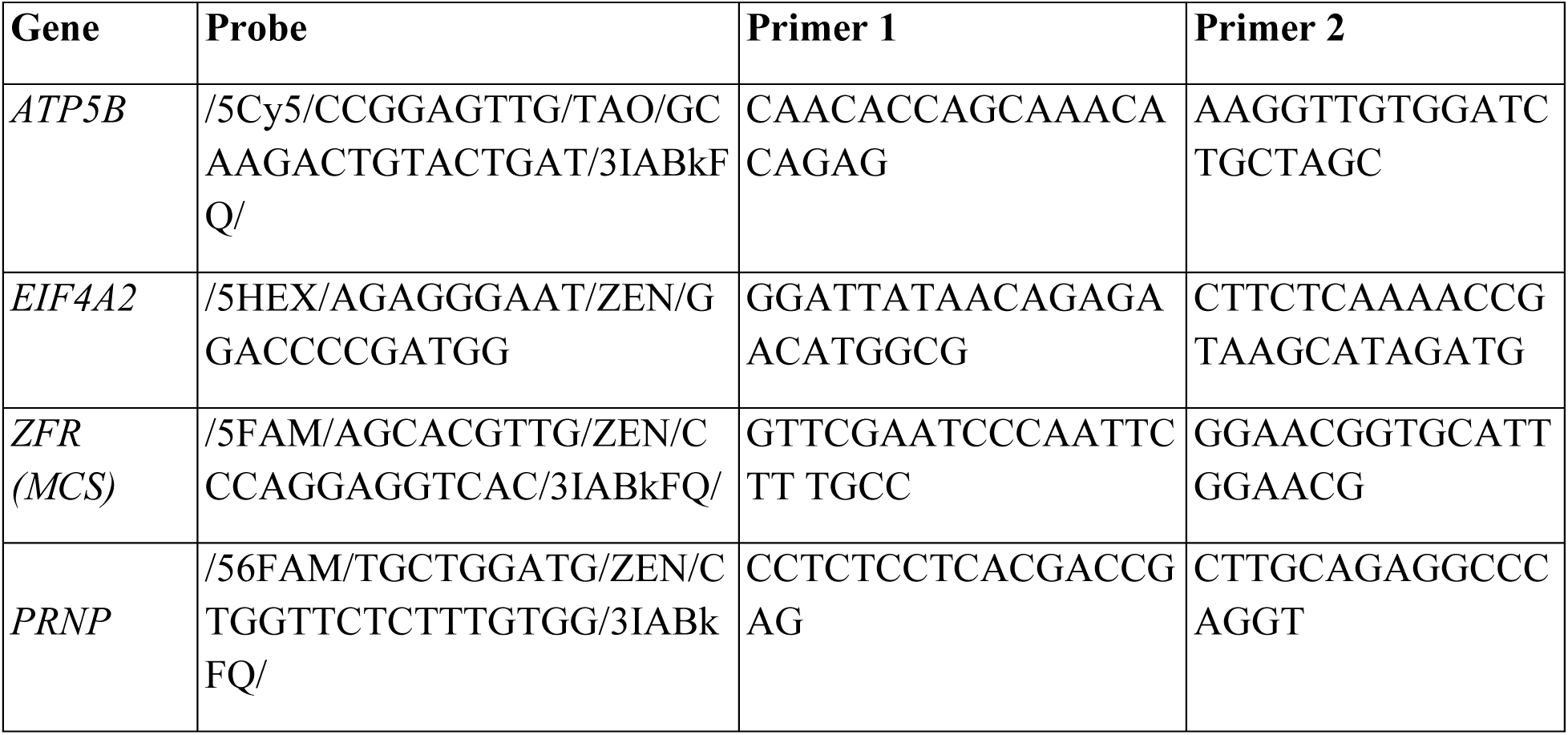
Taqman RT-qPCR Probes (NHP)

RT-qPCR data were normalized to the mean of three housekeeping genes in mouse (*Atp5b*, *Eif4a2*, and *Gapdh*) and NHP (*ATP5B* and *EIF4A2*). For all endogenous transcripts, normalized gene expression levels were scaled to the mean of the vehicle-treated animals. Absolute transcripts (copies/ ng RNA input) of ZFR for each sample were calculated using the following equation:

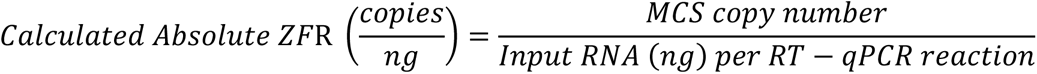

Data analysis and normalization were performed using Bio-Rad CFX Maestro program. Data were graphed using Prism 9 software (GraphPad Software; San Diego, CA). The data were analyzed using ANOVA. For statistical analyses, one-way ANOVA with Dunnett’s multiple comparisons test was conducted. An effect was considered significant if p < 0.05.

### Single-cell multiplexing ISH/IHC protocol (RNAscope) - Mouse Studies

Sections of 4 µm mouse brain slices were hybridized with a combination of two probes, panZFN-KRAB-C2 (ACD, #851651-C2) and Mm-PrnP-C1 (ACD, #476611) for detection of ZFR and *Prnp* mRNA, respectively. RNAscope® Multiplex Fluorescent Reagent Kit v2 (ACD, #323110) was used for ISH color development. Following ISH, brain slices were stained with antibodies against an anti-NeuN (Millipore, #ABN90P) or anti-S100β (Abcam, #ab52642) antibody, to label putative neurons or astrocytes, respectively. Secondary antibodies used were goat anti-guinea pig (Abcam, #ab175758) and goat anti-rabbit (Sigma, #SAB4608184). DAPI (Sigma, #D9542) staining was additionally performed to label all cellular nuclei. Images were acquired with the slide scanner Axio Scan.Z1 from ZEISS using a 20x Plan-apochromat objective (0.8 M27). Regions of interest (ROIs; thalamus, hippocampus, cortex, and caudate/putamen [CP]) were manually outlined in each image using the software Zen (v.2.3, Carl Zeiss Microscopy GmbH). Image analysis scripts for quantification of ZFR-mediated *Prnp* repression were developed using Acapella Studio 5.1 (PerkinElmer Inc.) and the integrated Acapella batch analysis as part of the Columbus system (v 2.9.1, PerkinElmer Inc.). Briefly, in each tissue section, the staining intensity per channel was normalized to the background. Individual cells within tissue sections were identified using the DAPI signal. Detection of ZFR and *Prnp* expression as individual spot was calculated using the SER Spot Texture Filter of Acapella 5.1 with a spot texture filter scale of 2 px for DAPI and 1 px for ZFR and *Prnp*. Spots were segmented as objects with a texture fluorescence signal above 0.1 for DAPI and 0.01 for ZFR and *Prnp*. Objects smaller than 7 px for DAPI and smaller than 5 px for ZFR and *Prnp* were excluded from the analysis. Furthermore, only objects were included that had a mean fluorescence intensity higher than the global mean intensity by 5-fold for DAPI and higher than 2-fold for ZFP and *Prnp*. Finally, the mean spot intensity had to be higher compared to the local surrounding by 2-fold for DAPI, by 1.7-fold for ZFR and by 1.5-fold for *Prnp*.

### Single-cell multiplexing ISH/IHC protocol (RNAscope) - NHP Study

Sections of 4µm nonhuman primate brain slices were hybridized with probes to detect *PRNP* (Mfa-PRNP-C1 # 1307661-C1, ACD) and the hZFR1 sequence (Custom probe design, ACD). Experimental procedures of ISH/IH protocol and imaging acquisition were the same as used in mouse study. Detection of ZFR and *PRNP* expression as individual spot was calculated using the SER Spot Texture Filter of Acapella 5.1 with a spot texture filter scale of 1 px. Spots were segmented as objects with a texture fluorescence signal above 0.1 for ZFR and 0.15 for the *PRNP* channel. Objects smaller than 4 px were excluded from the analysis. Furthermore, only objects with a mean fluorescence intensity higher than 2-fold the global mean intensity were included. Finally, the mean spot intensity had to be higher compared to the local surrounding by 2-fold for ZFR and by 1.4-fold for PRNP.

#### Quantification of transduced Neun+ cells – NHP study

To quantify transduced neurons, we performed image segmentation on both the NeuN and GFP channels using fine-tuned models based on Cellpose’s cyto3 model. For each channel, we manually annotated four random sites from the brain’s pons region, splitting the 2048×2048 image from each site into 512×512 subimages, which were then randomly divided into training and validation sets. Fine-tuning was performed with the following parameters: n_epochs=500, weight_decay=1e-4, learning_rate=0.01, and SGD=false for both the NeuN and GFP channels, resulting in fine-tuned NeuN and GFP models. These models were subsequently applied to other sites and brain regions, with manual corrections made to the resulting masks when necessary. The NeuN and GFP masks were then combined to produce double-positive masks requiring at least 40% overlap, which were subsequently counted to quantify transduced Neun+ cells in each brain region and site.

### RNAseq Analysis

#### Single nucleus sequencing

Brain cortical tissue was collected from the promoter comparison study for single nucleus RNA-seq (snRNA-seq) analysis. Nuclei were purified and 10X single nucleus sequencing performed as described (Mortberg et al. 2023). Briefly, single nucleus suspensions were generated following the protocol described by (Kamath et al. 2022) using trituration with a pipette in a buffer of Kollidon VA64, Triton X-100, bovine serum albumin, and RNase inhibitor. Suspensions were extruded through a 26-gauge needle, washed, pelleted, and strained. Nuclei were sorted for positive DAPI signal on a Sony SH800 or MA900 cell sorter with a 70 µm chip using a 405 nm excitation laser and a 425 - 475 nm filter for light collection. Nuclei counts were obtained by hand tally counter with a Fuchs-Rosenthal C-Chip hemocytometer. Volume was measured to target 17,000 nuclei and submitted to the Broad Institute’s Genomics Platform, for 10X library construction (3’ V3.1 NextGEM with Dual Indexing) according to manufacturer instructions [10X Genomics USER GUIDE: chromium Next GEM Single Cell 3′ Reagent Kits v3.1 (Dual Index). 2022; CG000315 Rev E.] 100 cycles of sequencing were performed on an Illumina Novaseq 6000 S2.

#### Single-nucleus sequencing data processing, filtering, and cell type assignment

Single nucleus RNA-sequencing libraries were sequenced on a NovaSeq S2. Read quality control, UMI counting, barcode counting, and alignment to a custom genome (mm10 GENCODE vM23/Ensembl 98 from 10x Genomics and the mZFR1 sequence) were performed using the “cellranger 7.0.0” pipeline. All downstream analysis were performed using the Seurat packages in R for single-cell RNAseq analysis. All samples were filtered on the following measures: number of unique genes detected, total number of molecules detected, and level of expression of mitochondrial reads before integrating the datasets. We removed cells in which greater than 5% mitochondrial reads, less than 200 feature counts, or greater than 6000 feature counts were detected. For cell type assignment, data were normalized using the Seurat normalization function which divides the feature counts by total counts for each cell, then multiplies the resulting fraction by a scale factor of 1.00e+4, and, lastly, log-normalizes this value. Dimensionality reduction was performed using Seurat packages and utilizing principal components 1 through 10 in the reduction. Clusters were determined at a resolution of 0.7. The identity of each cluster was validated by evaluation of gene expression within each cluster of cell-type specific markers for the cell-types of interest.

#### Binomial Modeling

For this experiment the same statistical approach as (Mortberg et al. 2023) was employed. A negative binomial fit using the MASS package in R was used: glm.nb(prnp_umis ∼ celltype + celltype:treatment + offset(log(total_umis))). The model returns coefficients in natural logarithm space, so for treated conditions, the coefficient for each cell type-treatment interaction term coefficient was then exponentiated to yield the mean the residual target RNA, while the 95% confidence interval was defined as that mean ±1.96 of the model’s standard error. The point estimate for each animal was computed by adding the model’s residual to the cell type-treatment coefficient, and then exponentiating. ZF+ cells were simply defined as those with at least 1 UMI of the ZFR transcript added to the custom reference.

### PrP Quantification in Mouse Brain by ELISA

PrP was quantified by ELISA in frozen whole hemispheres (Mortberg et al. 2022). Left brain hemispheres were homogenized at 10% wt/vol in 0.2% wt/vol CHAPS (Sigma; C9426). The 96-well plate assay uses anti-PrP antibody EP1802Y (Abcam; ab52604) for capture, and biotinylated anti-PrP antibody 8H4 (Abcam; ab61409) for detection, followed by streptavidin-HRP (Thermo Fisher Scientific, 21130) and TMB (Cell Signaling, 7004P4). Recombinant mouse PrP protein (Broad Institute; Lot No. MoPrP16 or MoPrP23-231) was used to generate a standard curve (Reidenbach et al. 2020). Data presented are percentages of residual PrP normalized to the mean of vehicle controls.

### PrP Quantification in Mouse CSF by Electrochemiluminescence Assay

Samples were added to the collection buffer immediately. Mouse CSF PrP was quantified by electrochemiluminescence assay using Meso Scale Discovery (MSD) technology. The assay uses biotinylated anti-prion antibody 6D11 (Biolegend, 808002) as capture antibody and sulfo-tag conjugated anti-prion antibody 8H4 (Abcam, ab61409) as detection antibody. Capture antibody was loaded to 96-well small spot streptavidin plates (MSD, L45SA-1). After the incubation of detection antibody in the plates, MSD gold read buffer B (MSD, R60AM) was added to the plates to generate electrochemiluminescence signals. Signals were read using MESO SECTOR S 600MM. Recombinant mouse PrP was generated from Impact Biologicals using an E.coli expression system and used as the calibrator for standard curve preparation. Final PrP concentrations were calculated based on the dilution factors of CSF samples in the buffers. Data presented are percentages of residual PrP normalized to the mean of vehicle controls.

### AAV packaging and production of ZFRs

Recombinant adeno-associated viral vectors (rAAV) were generated by the triple transfection method. Three plasmids – (i) an AAV Helper plasmid containing the Rep and Cap genes, (ii) an Adenovirus Helper plasmid containing the adenovirus helper genes, and (iii) a transgene plasmid containing the sequence to be packaged flanked by AAV2 inverted terminal repeats were transfected into the HEK293 cells using calcium phosphate. Harvested cells were lysed three days after transfection and the rAAV was precipitated using polyethylene glycol. Virus produced was further purified by ultracentrifugation overnight on a cesium chloride gradient. The virus was formulated by dialysis and then filter-sterilized. After adjusting the titer (vector genomes/mL; vg/mL) of all AAV batches by dilution with PBS + 0.001% Pluronic F-68, the AAVs were aliquoted to single use doses and stored at -80°C until use. No samples were frozen after they were thawed.

### Statistical Analysis

Statistical analyses for data from RT-qPCR, ELISA, and single cell ISH/IHC were conducted with GraphPad Prism Version 10. Analyses at Broad were conducted using custom scripts in R 4.2.0. Binomial modeling for the single nucleus sequencing data is described above. Survival curves utilized the R *survival* package and differences in survival were assessed using 2-sided log-rank test. Animal body weights were normalized to individual baseline and then averaged by treatment group. Shaded areas in plasma NfL and normalized weights plots represent standard deviation of the mean for each treatment group. For nesting and activity scores, a Gaussian-smoothed average is shown, produced using the R *smoother* package with a window of 25 days. P values less than 0.05 were considered nominally significant. Raw data and source code sufficient to reproduce our analyses will be made publicly available at github.com/ericminikel/zfr and single nucleus sequencing data will be deposited at singlecell.broadinstitute.org.

## Acknowledgements

We thank S. Bhardwaj, T. Chen, S. Girdhar, A. Goodwin, H. Tran and C. Wu for AAV production. We thank A. Hatami, M. Glynn, M. Johnson and J. Stallons for mouse study support. We thank L. Andrews, J. Cifrese, S. Frias, J. Hodges, G. Houlihan, K. Kim, J. Lee, M. Mendel, A. Nanjaraj, A. Olin, E. Ponce, M. Samie, Y. Santiago and A. Young for NHP study tissue analysis. We thank M. Lanier for sample testing. We thank M. Chen and M. Tian for assisting with statistical analysis. We thank C. Gasper, K. Lewis, S. Mueller and J. Perez for operations support. We thank M. Jensen, S. Mueller and E. Paysinger-Hill for project management. We thank S. Schreiber and A. Woolfson for initial program support. We thank N. Dubois-Stringfellow, J. Fontenot and A. Macrae for program support. We thank G. Atkinson and A. Resch for program oversight. We thank C. Courme for illustration support. We thank G. Brasnjo, C. Cocciardo, G. Davis, N. Dubois-Stringfellow, V. Headley, A. Macrae, E. McNeil, P. Ramsey and L. Wilkie for manuscript review. We thank G. Atkinson for helpful discussions.

---

**Supplementary Figure 1.**
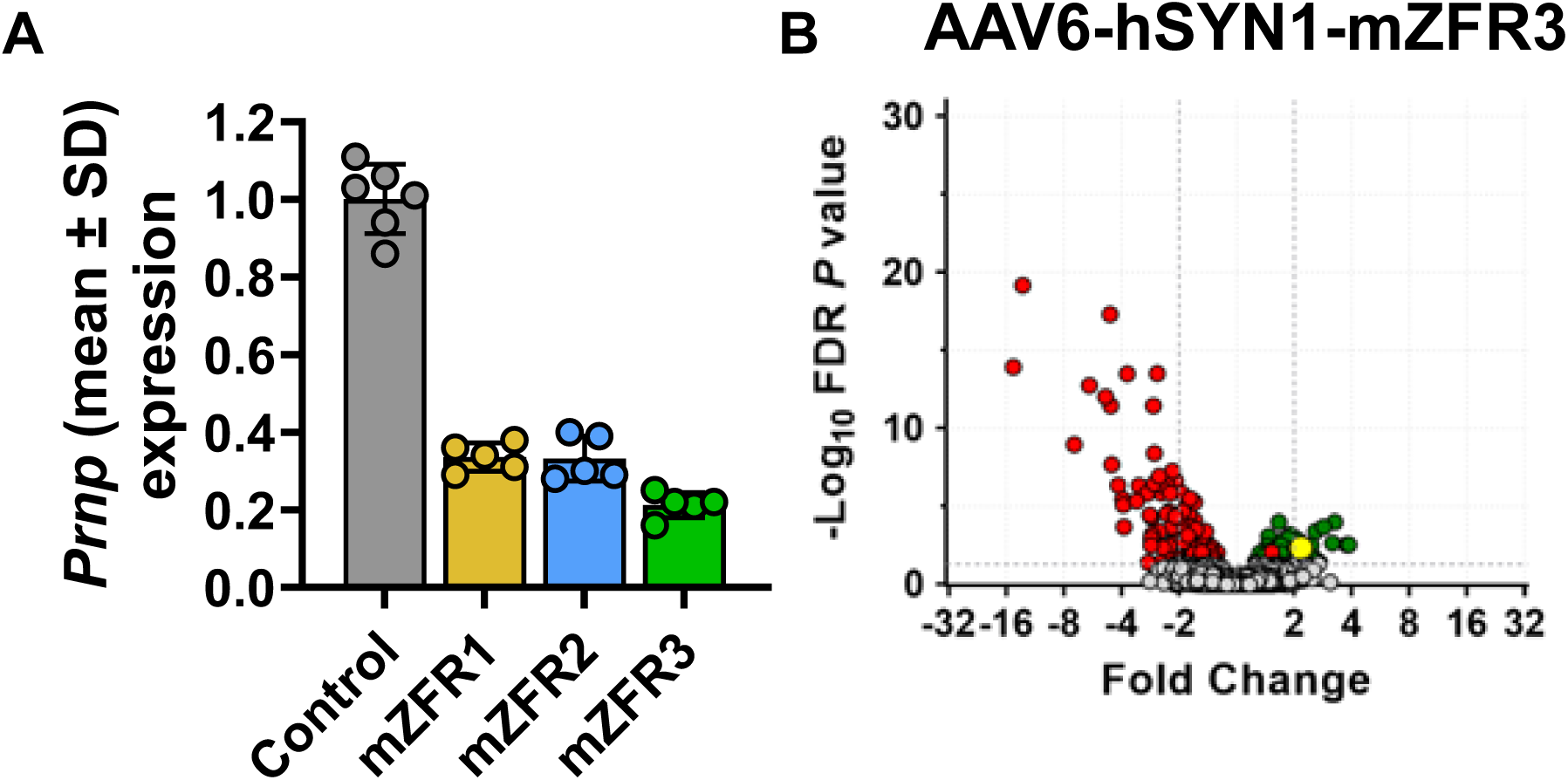
Example of a mZFR with potent target gene repression with many differential regulated genes. **(A)** Corresponding RT-qPCR assessment of *Prnp* gene expression from mouse cortical neurons transduced with AAV6-mZFRs at 3E+3 MOI for the affymetrix analysis shown in Figure 1C and in Supplemental Figure 1B. Mean ± SD. n=5-6 biological replicates for per treatment. **(B)** Affymetrix array analysis for a less specific ZFR (mZFR3) showed many up-(green circle) and down-regulated (red circle) transcripts (significance threshold FDR *P* <0.05), confirming the sensitivity of the assay to detect dysregulated genes.

**Supplementary Figure 2.**
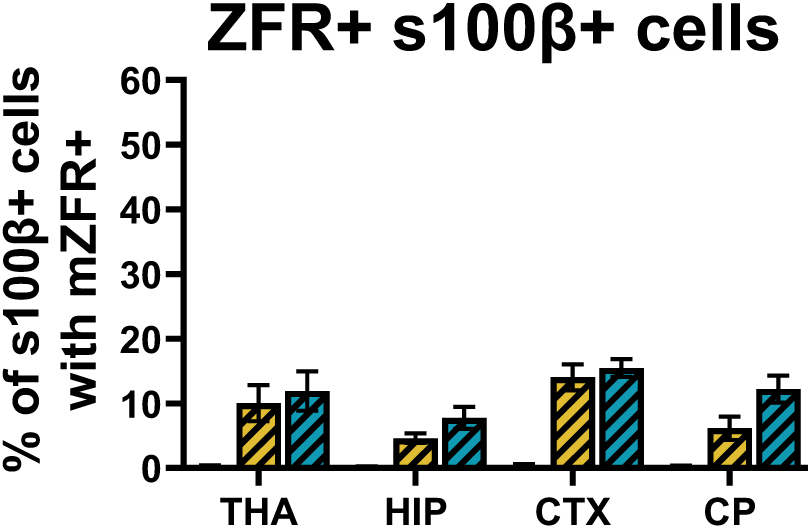
Percentage of s100β+ cells with ZFR expression. Percentage of ZFR+ s100β+ cells (mean ± SD) in the thalamus (THA), hippocampus (HIP), cortex (CTX), and caudate/putamen (CP). Depending on the brain region, the percentage ranged from 4.6-14.0% for the hSYN1 group (yellow diagonal bars) and 7.8-15.5% for the GfaABC1D group (green diagonal bars). Due to the poor staining quality, data was not quantified for the CMV group.

**Supplementary Figure 3.**
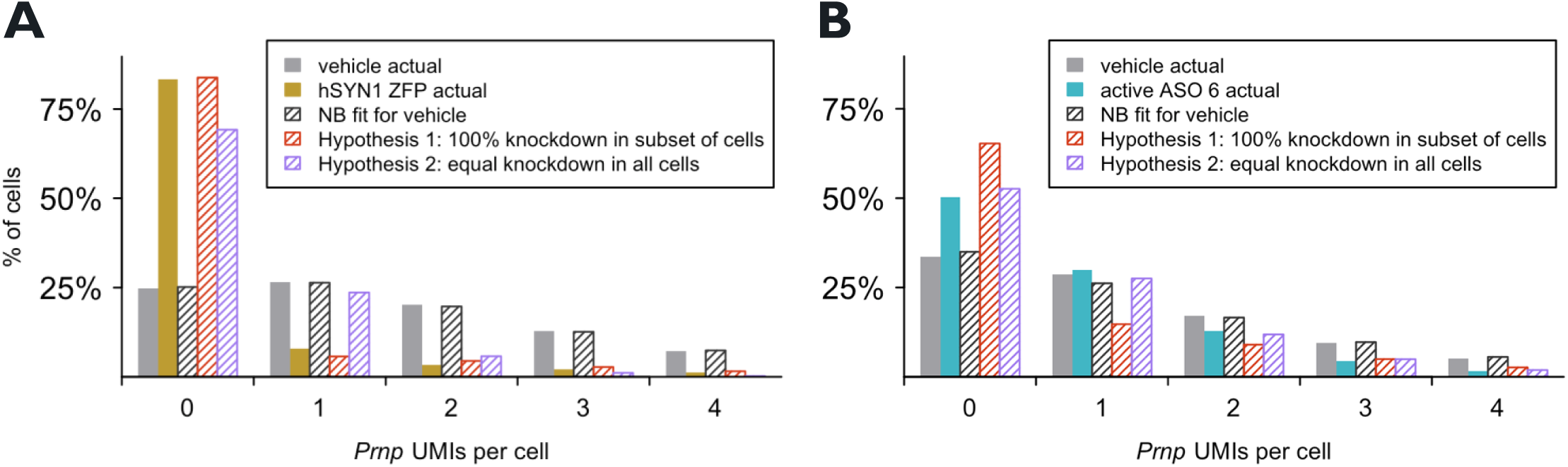
Histograms of *Prnp* counts in single cortical neurons. Histograms of *Prnp* in single cortical neurons (excitatory and inhibitory) for ZFR **(A)** and ASO **(B)**. The data in (B) is the re-analysis of the data from Mortberg & Gentile (2023) using animals that received 500 µg active ASO 6 and harvested at 2 weeks post-dose. Data is shown on the same y-axis scale as the ZFR data to allow comparison. (Note that Figure 2F from Mortberg & Gentile (2023) is astrocytes at 12 weeks post-dose and shown on a different y-axis scale).

**Supplementary Figure 4.**
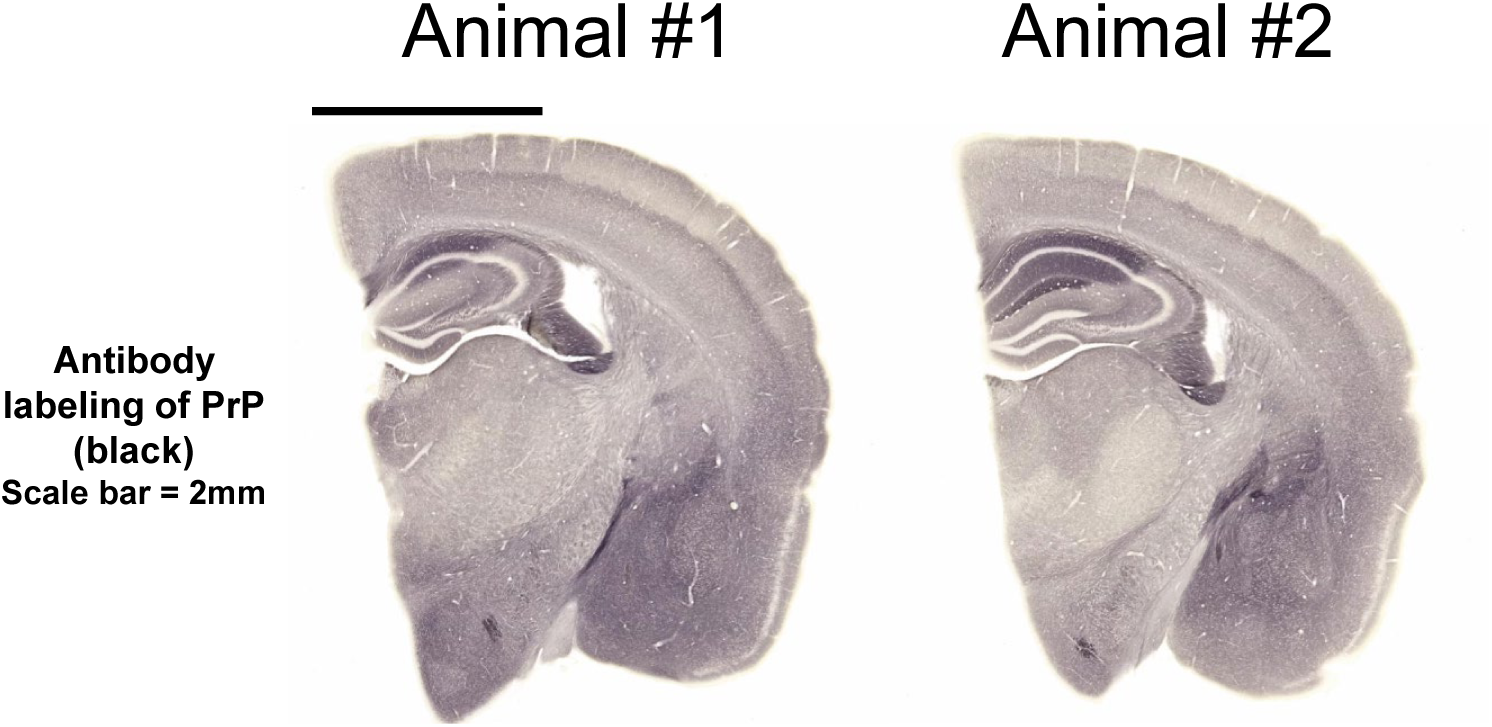
PrP protein expression from uninoculated and untreated mouse brain. Representative images with staining for PrP expression on brain sections from uninoculated, untreated mice that survived to the end of Survival Study A (500 dpi).

**Supplementary Figure 5.1.**
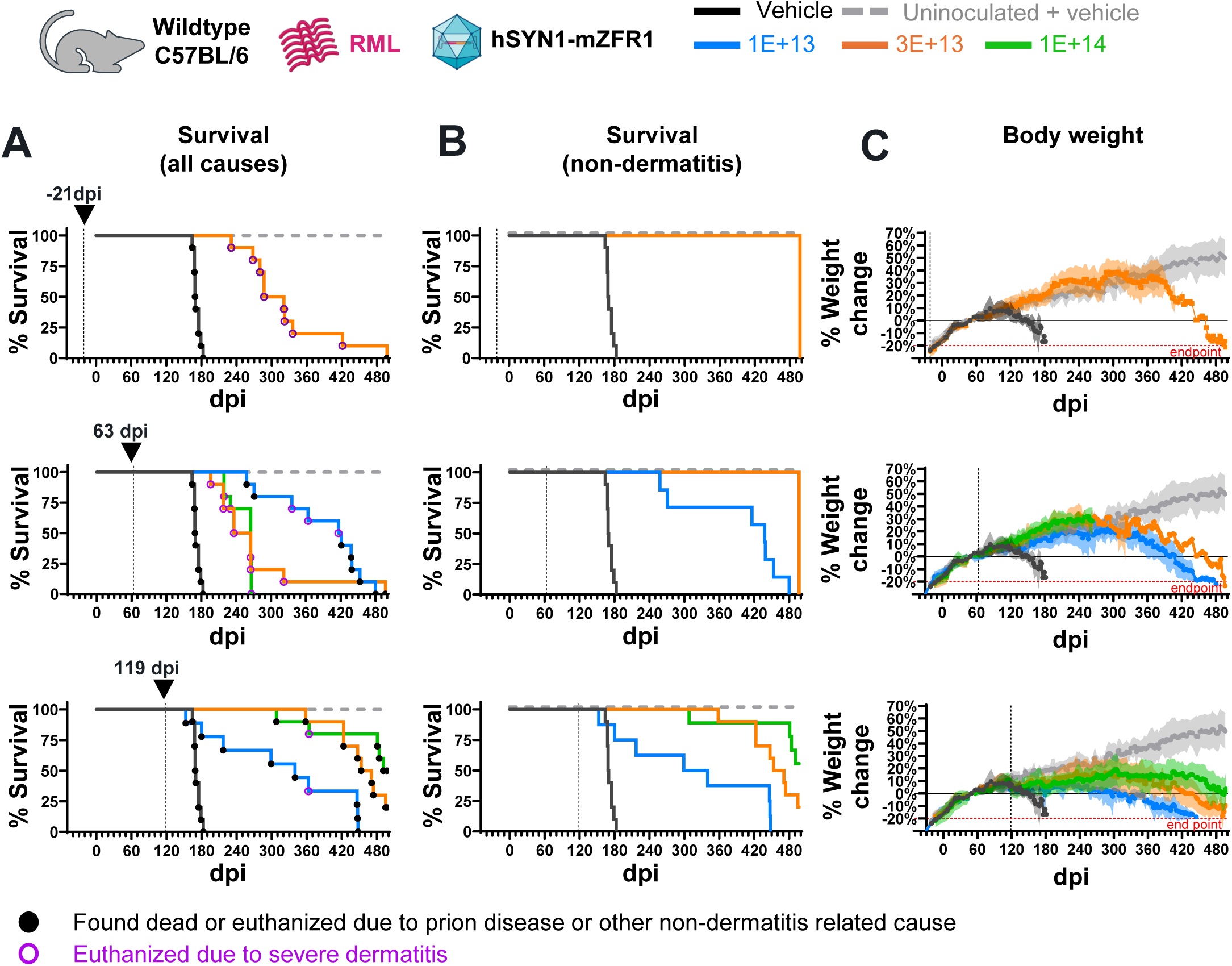
Treatment of hSYN1-mZFR1 extends survival at different stages of prion disease in mice. **(A)** Survival curves representing all-cause mortality for test groups that received hSYN1-mZFR1 at -21 (top), 63 (middle), and 119 (bottom) dpi. Each plot shows vehicle (black line), 1E+13 vg/kg low dose (blue line), 3E+13 vg/kg mid dose (orange line), and 1E+14 vg/kg high dose (green line) groups (n=10/group, except n=9 in low dose 119 dpi group). The dashed grey line represents vehicle-treated uninoculated mice (n=5). Animals that were found dead or were euthanized due to non-dermatitis endpoint are labeled with black closed circles, while animals euthanized due to severe dermatitis are labeled with purple open circles. **(B)** Survival curves plotted with animals euthanized for intercurrent severe dermatitis censored (9 animals in the mid dose -21 dpi group; 3, 9, and 10 animals in the 63 dpi low, mid, and high dose groups, respectively; and 1 animal in each of the 119 dpi low and high dose groups). **(C)** Percent body weight change relative to the individual baseline weight at 49 dpi plotted as mean ± SD (solid line ± shaded area).

**Supplementary Figure 5.2.**
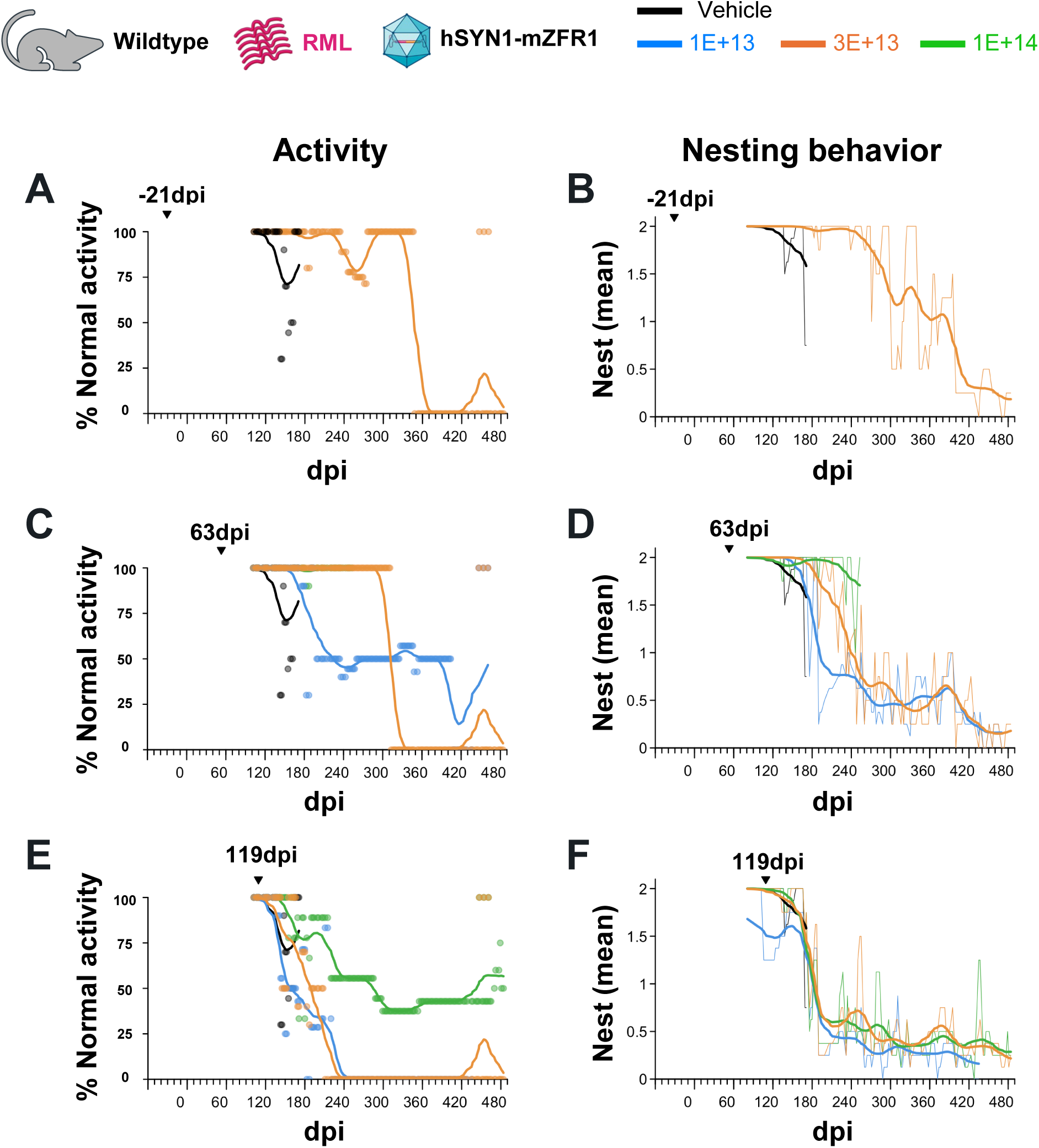
Effect of hSYN1-mZFR1 treatment on animal behavior at pre-symptomatic, pathological and symptomatic timepoints. Animals were inoculated with RML prions on day 0, and intravenously administered vehicle or hSYN1-mZFR1 at different doses on either -21 dpi **(A, B)**, 63 dpi **(C, D)**, or 119 dpi **(E, F)**. Group size 10 animals per group for vehicle (black), 1E+13 vg/kg (blue), 3E+13 vg/kg (orange), and 1E+14 vg/kg (green). Animals in an additional control group received vehicle but no RML prions (grey, n = 5). **(A, C, E)** Dots indicate the proportion of animals in a given group that showed normal activity plotted against days post-RML-inoculation, such that 100% means all animals showing normal activity and 0% means no animals with normal activity. The curves are smoothed means of the points (Gaussian smoother with window of 25 days and with tails included, using the function smth from the R package smoother). **(B, D, F)** Nest-building activity. Animals were housed 5 per cage and nesting behavior was evaluated on a per cage basis for nestlet and Enviro-dri utilization, see Methods. Thin lines show the mean score across cages within each group at any individual observation session, while the thick lines show smoothed means (Gaussian smoother with window of 25 days and with tails included, using the function smth from the R package smoother).

**Supplementary Figure 5.3.**
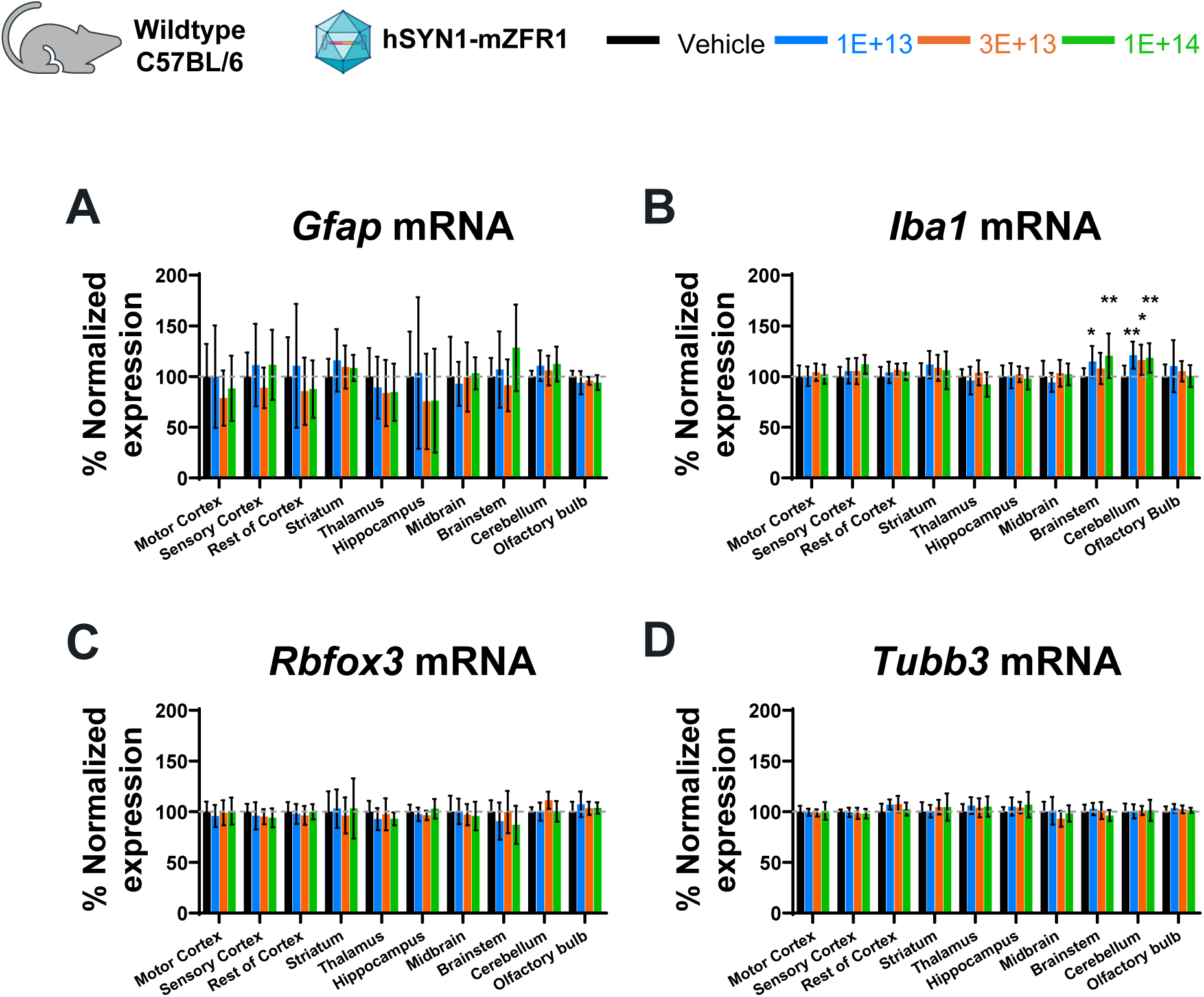
Impact of hSYN1-mZFR1 administration on neuronal and glial markers in mouse brain. **(A-D)** RT-qPCR results from different brain regions for each treatment group corresponding to Figure 5D. Normalized expression values were scaled to the mean of the vehicle-treated group for each transcript (Mean ± SD). **(A)** *Gfap*, Glial fibrillary acidic protein; **(B)** *Iba1*, ionized calcium-binding adapter molecule **(C)** *Rbfox3*, protein marker for neuronal marker; **(D)** Tubb3, neuronal specific tubulin (*Tubb3*). Two-way ANOVA analysis using Dunnett’s multiple comparisons test. No statistically significant variation was found in hSYN1-mZFR1 treated group at any dose when compared to the vehicle-treated group, except the *Iba1* expression from brainstem of 1E+13 group (115%, p = 0.04) and 1E+14 group (120%, p = 0.003) and cerebellum of 1E+13 group (121%, p = 0.002), 3E+13 group (116%, p = 0.02) and 1E+14 group (118%, p = 0.007).

**Supplementary Figure 6.**
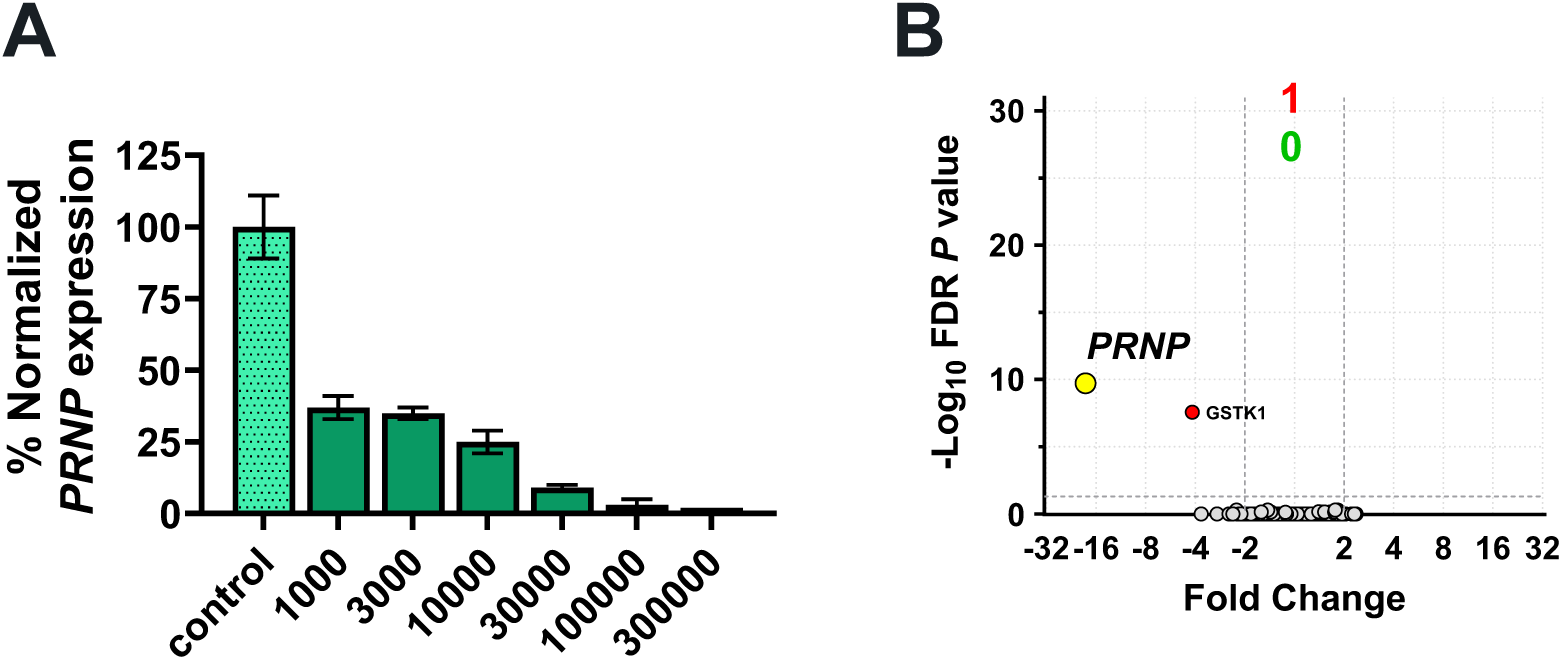
Potent and specific *PRNP* repression by hZFR designed to target the human *PRNP* gene. **(A)** Normalized *PRNP* mRNA expression values from human iPSC-derived neurons (iGABA) 19 days after transduction with AAV6-hSYN1-hZFR at six different MOIs (x-axis). Values were scaled to control treatment (mean ± SD) **(B)** Affymetrix analysis for iGABA neurons 19 days after transduction with AAV6-hSYN1-hZFR at 1E+5 MOI (n=6 biological replicates). A yellow circle denotes the human *PRNP* gene.

